# Rad53/Chk2 couples mitochondrial outputs and ROS with G2/M cell cycle progression

**DOI:** 10.64898/2026.10.14.759190

**Authors:** Swagata Adhikary, Ritu Gupta, Vineeth Vengayil, Sunil Laxman

## Abstract

Cell-cycle progression requires coordination with cellular metabolism, yet how checkpoint kinases contribute to this coordination during normal growth remains poorly understood. Here, we uncover an unexpected role for the checkpoint kinase Rad53/Chk2 in coupling mitochondrial and redox homeostasis to G2/M progression. In prototrophic *S. cerevisiae,* Rad53 loss (*rad53Δ*) disrupts cell-cycle progression and causes cells to accumulate in G2/M, alongside extensive metabolic remodelling with increased tricarboxylic acid (TCA) cycle flux. Mitochondrial proteomics reveals altered relative abundances of individual electron transport chain proteins, and mitochondrial membrane potential increases without increased oxygen consumption or ATP generation. This altered mitochondrial state is associated with elevated reactive oxygen species (ROS) and oxidative stress. Genetic analyses establish that mitochondrial phenotypes require Rad53 kinase activity and are distinct from the loss of SML1. Antioxidant treatment rescues cell-division, budding defects and the accumulation of abnormally sized cells, establishing ROS as a primary cause of cell-cycle defects. Thus, Rad53 couples mitochondrial and redox homeostasis to G2/M progression during normal growth.

## Introduction

The cell cycle stages are tightly regulated by cell cycle checkpoints, which also monitor the fidelity of processes such as DNA replication, DNA integrity, and chromosome segregation^1,2^. These checkpoints are coupled with signaling outputs by cyclin-dependent kinases (CDKs), ubiquitin ligases, phosphatases, and checkpoint kinases^3,4^. It is now clear that cell cycle progression is closely coupled to the cellular metabolic state^5–7^. Orderly progression through the cell cycle checkpoints occurs only when the metabolic requirements of each phase of the cell cycle are satisfied^8–10^. There are distinct metabolic events associated with each cell cycle phase. For example, high acetyl CoA levels drive entry into the G1 phase, SAM and methionine are essential for G1/S progression^4,8,9^, while the S-phase requires a reduced cytoplasmic environment to maintain genome integrity^11,12^, and sustained anabolism is required for G2/M progression^5^. Importantly, any deviations from these metabolic and redox states can influence the cell cycle states. In particular, disrupting redox homeostasis and accumulating high reactive oxidative species (ROS) activate checkpoint pathways to delay cell cycle progression^11,13^. Studies in budding yeast revealed that periodic changes in metabolic/redox state are coordinated with cell division^5,14^. Interestingly, the disruption of oxidative stress defense pathways impairs this timely coordination of metabolism and cell division by invoking checkpoint pathways ^11,15^. Checkpoint pathways therefore provide an important mechanism through which cells can respond to metabolic and redox perturbations and adjust cell cycle progression accordingly. While these requirements are clear, how checkpoint proteins might couple metabolic-redox homeostasis to cell cycle progression remain poorly understood.

Rad53, the *S. cerevisiae* homolog of mammalian Chk2, is an extensively studied DNA damage checkpoint kinase, which maintains genome integrity in cells encountering exogenous DNA damage inducing agents^16,17^. Rad53 is activated by hyperphosphorylation upon DNA damage by damaging agents, and then regulates downstream processes leading to cell cycle arrest^18,19^. Upon DNA damage, Rad53 inhibits Sml1, which is an inhibitor of ribonucleotide reductase (Rnr1). This action of Rad53 activates Rnr1, leading to increased dNTP synthesis to allow DNA repair^20–22^. While these are well-established roles during DNA damage, the functions of RAD53 during homeostatic growth in the absence of damaging agents, remain unclear. Historically, studying these DNA damage-independent functions of Rad53 has been challenging since in the commonly used, multiple auxotrophic laboratory strains of yeast, the deletion of Rad53 is lethal^20,23^. Instead, historically studies in laboratory auxotrophic strains relied on viable, secondary suppressor mutations of Sml1 (*sml1Δ*) in a *rad53Δ* background, where the DNA-damage specific roles of Rad53 can be specifically studied ^21,23^. However, recent studies suggest possible roles for Rad53 managing metabolic stress^24,25^, or its activation independent of the DNA damage axis^26^, suggesting broader homeostatic functions of Rad53.

Here, we discover a central role for Rad53 in restraining mitochondrial outputs, to enable normal cell cycle progression. Rad53 knock-out cells are viable in fully prototrophic yeast cells. The loss of Rad53 in these cells leads to delayed cell cycle progression and G2/M arrest, alongside a distinct metabolic rewiring. Quantitative metabolite and flux experiments in *rad53Δ* cells reveal altered glycolytic and increased TCA cycle flux. Proteomic and functional analyses of mitochondria from *rad53Δ* cells revealed an altered mitochondrial composition, without increased oxygen consumption or mitochondrial ATP production, leading to severe oxidative stress. Rescuing ROS in cells in *rad53Δ* cells restores normal cell cycle progression. These findings reveal an essential, homeostatic role for Rad53 in restraining mitochondrial outputs and ROS to allow normal G2/M progression, suggesting that Rad53 checkpoint kinase functions as a metabolic gatekeeper for normal cell cycle progression and growth.

## Results

### Loss of Rad53 kinase results in G2/M arrest, delayed division and growth

Rad53 is a Ser/Thr checkpoint kinase, well studied for its role in mediating responses to acute DNA damage. Upon treatment with DNA damaging agents (such as methyl methane sulfate (MMS)), Rad53 is hyperactivated and drives the DNA damage repair pathway (Fig 1A). Contrastingly, functions for this kinase during homeostatic cell proliferation in absence of exogenous DNA damaging conditions remain poorly understood (Fig1A). We qualitatively assessed the relative protein amounts and phosphorylation status of Rad53 in cells during growth in standard high glucose media (YPD), and upon inducing DNA damage. Upon treatment with DNA damaging agents such as methyl malonyl sulfate (MMS), Rad53 is hyperphosphorylated – observed as reduced-mobility bands on SDS-page gels (Fig 1B). In contrast, in yeast cells without any exogenous DNA damaging conditions, the Rad53 protein remained constitutively present but without hyperphosphorylation (Fig 1B). We also assessed relative Rad53 protein amounts and phosphorylation across cell cycle phases, following release from α – factor mediated G1 arrest. Rad53 electrophoretic mobility remained ∼ unchanged across the cell cycle phases (Fig S1A). Collectively, we observe constitutive steady levels of Rad53 protein, and where the damage-induced hyperphosphorylation was not present during normal cell proliferation.

**Fig 1.**
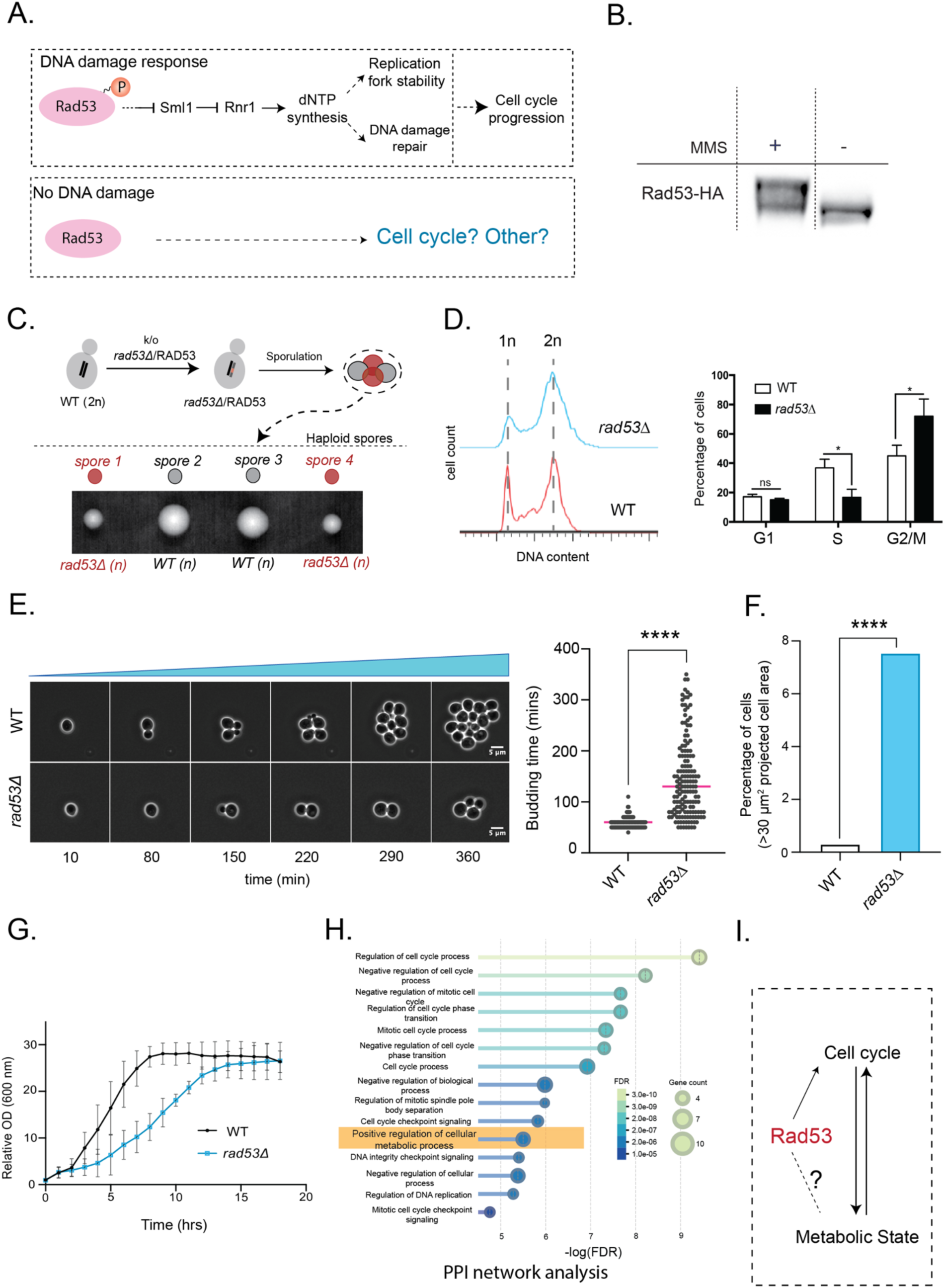
Rad53 regulates G2/M progression in the absence of exogenous DNA damage. **A.** A schematic illustrating known Rad53/Chk2 functions in the DNA-Damage response pathway, while other functions of Rad53 during cell division are unclear. **B.** Assessment of Rad53 electrophoretic mobility in the presence or absence of DNA-damage inducing agent, MMS. A HA-epitope tag was introduced to the endogenous *RAD53* locus. Electrophoretic mobility on SDS-PAGE gels as detected by Western Blotting is shown. MMS treatment results in hyperphosphorylated Rad53 is observed as reduced electrophoretic mobility bands on SDS-PAGE gels, vs faster electrophoretic mobility during grown in standard media without MMS. Also see Fig S1A for Rad53 electrophoretic mobility across cell cycle phases. **C.** Generation of viable *rad53Δ* cells in fully prototrophic yeast. The schematic and data show tetrads of heterozygous *rad53Δ*/RAD53 diploid yeast, followed by sporulation and dissection diploids yielding two wild-type and two viable *rad53Δ* haploid spores per tetrad. Also see Fig S1B for genotype analysis. **D.** (Left) Analysis of DNA content by flow-cytometry. Flow analysis indicates that *rad53Δ* cells have delayed cell-cycle progression, along with a significantly increased proportion of cells accumulating in the G2/M phase. (Right) Quantification of the flow-cytometry data shown. Data are displayed as means ± SD, n = 3. *p<0.05, **p<0.01, ***p<0.001, ****p<0.0001, student’s T test. **E.** (Left) Representative time-lapse live cell images of WT and *rad53Δ* cells in standard media, acquired continuously over 360 minutes of growth. Time (min) is indicated below each frame. Scale bar - 5 μm. (Right) *rad53Δ* cells exhibited a prolonged budding time, with a median of 130 min compared with 60 min for WT cells (WT *n* = 120; *rad53Δ n* = 150, three independent experiments). Individual data points are shown, with the median indicated. ****p<0.0001, two-tailed Mann–Whitney U test. Also see Fig S1C for interquartile range (IQR) of budding time and budding time distribution. Movie S1 and S2 show movies of cell division of the respective WT and *rad53Δ* cells over time. **F.** Fraction of total cell population that is extremely enlarged (>30 μm²) in WT and *rad53Δ* cells (also see Methods for analysis details). *rad53Δ* cells exhibited a significant increase in the proportion of enlarged cells compared with WT. Statistical significance was determined using Fisher’s exact test (****p<0.0001, odds ratio = 27.6; WT, *n* = 600; *rad53Δ*, *n* = 700, pooled from three independent experiments). Also see Fig S1D for the fraction of cells in each bin. **G.** Liquid growth curves of WT and *rad53Δ* cells. *rad53Δ* cells exhibit slower growth and delayed biomass accumulation compared with WT cells. Data are presented as mean ± SD, n = 3 (per time point). Also see Fig S1E for growth on solid agar, and growth characterizations of distinct *rad53Δ* clones. Fig S1F shows the effect of DNA-damage inducing agents on WT and *rad53Δ* cells. **H.** Gene Ontology (GO) biological process enrichment analysis of the Rad53 protein– protein interaction network from the STRING database. Enriched biological processes are ranked by significance based on the false discovery rate (FDR). The most significantly enriched terms are shown, with ‘positive regulation of cellular metabolic process’ among the most significantly enriched terms. **I.** A hypothetical illustration suggesting a role of Rad53 in cell cycle regulation, alongside an unknown coupling with cellular metabolic state.

Since Rad53 has primarily been studied in the context of the DNA damage repair pathway, the roles of Rad53 in the absence of induced DNA-damage are poorly understood. Prior studies investigating Rad53 functions utilized auxotrophic laboratory strains of *S. cerevisiae*, which are deficient in several essential metabolic pathways and biosynthetic capabilities. The knock-out of Rad53 in these auxotrophic laboratory strains is lethal^20,23^. Therefore, studies of *rad53Δ* in these laboratory strains utilize an additional suppressor – the loss of Sml1 (downstream of Rad53 in the DNA damage response), which confers synthetic viability to *rad53Δ*^27^ in auxotrophic laboratory strains. *RAD53* knock-outs, in the absence of the suppressor *SML1* knock-out have not been studied, confounding the possibility of separating Rad53-dependent roles in DNA-damage from other outputs. In a robust, prototrophic, diploid strain (CEN.PK background), we deleted a single allele of *RAD53*. These diploid cells were then sporulated to tetrads. Surprisingly, this consistently resulted in the generation four viable haploid daughter spores from each tetrad, with two of them being *rad53Δ* (Fig 1C, Fig S1B). While *rad53Δ* spores resulted in colonies that were noticeably smaller than the wild-type, we serendipitously found that prototrophic *rad53Δ* cells are viable, without the additional introduction of a *sml1Δ* suppressor.

In these *rad53Δ* cells, we assessed growth and cell cycle progression under standard high-glucose media (YPD), in the absence of any exogenous DNA damaging agents. The populations of cells in distinct cell-cycle stages were first analyzed by flow cytometry, and this revealed a markedly increased accumulation of *rad53Δ* cells in G2/M phase compared to wild-type cells, suggesting delayed progression through G2/M (Fig 1D) in these cells.

To directly measure cell division over time, we next performed extensive time-lapse, live-cell microscopy-based analyses. WT and *rad53Δ* cells were continuously imaged for ∼360 min on YPD (2%) agar pads, and budding times were quantified for 120–150 cells from each background from independent experiments. *rad53Δ* cells exhibited a marked increase in budding time, with mean and median budding times of 145 and 130 mins respectively, compared to mean and median budding times of ∼60 mins in the WT cells. Analysis of the budding time distribution further revealed that WT cells exhibited a narrow distribution centered around 60-80 min, whereas *rad53Δ* cells displayed a much broader distribution spanning approximately 60–160 min, indicative of increased cell-to-cell variability in cell-cycle progression (Fig 1E; S1C; Movie S1, S2).

We further examined whether this delay was accompanied by changes in the cell size distributions of WT and *rad53Δ* cells. Cell size was analysed using the final frame of the time-lapse movies. We compared the two-dimensional projected cell areas obtained from Cell-ACDC segmentation masks of WT and *rad53Δ* cells. The segmentation masks were generated using the morphology reconstruction module implemented in Cell-ACDC, which employs the YeaZ 2.0 neural network-based segmentation and shape inference framework^28^ (see Methods for more details). Analysis of the projected cell area revealed the presence of a substantial, distinct subpopulation of enlarged *rad53Δ* cells. Approximately ∼6-7% of *rad53Δ* cells had projected areas >30 µm², indicating very large cells, compared with fewer than 0.3% of WT cells falling in this projected area range (Fig 1F, S1D).

Collectively, we observed that the loss of Rad53 results in delayed cell cycle progression, G2/M arrest, the accumulation of an enlarged-cell subpopulation, and heterogeneity in division time and cell size, indicating that Rad53 is required to gate G2/M cell cycle progression, in the absence of exogenous DNA damaging agents.

Next, we compared the overall biomass increase in wild-type and *rad53Δ* cells, both on solid agar as well as in liquid culture, in standard high-glucose media (Fig 1G and S1E). Here, we observed a delayed lag phase and slower biomass accumulation of *rad53Δ* cells compared to WT (Fig 1G and S1E), with these cells taking substantially longer than WT cells to eventually reach stationary phase and comparable biomass. As controls, we separately assessed the growth of *rad53Δ* cells in presence of exogenous DNA damage (MMS, HU). *rad53Δ* cells expectedly exhibit severe growth defects (and hypersensitivity) upon DNA damage, consistent with the established role of Rad53 in the DNA damage response (Fig S1F). Together, these findings establish an essential role for Rad53 as a checkpoint for normal cell cycle progression and growth, in the absence of exogenous DNA damaging agents.

This role for Rad53 in normal cell cycle progression is understudied in current literature. In order to further address possible Rad53 roles in cell proliferation we mined existing interaction information for Rad53, in order to integratively assess overall processes regulated by Rad53. Using Gene Ontology (GO) Biological Process enrichment analysis of the protein–protein interaction network for Rad53 from the STRING DATABASE, we identified highly significant biological processes among the top 15 enriched terms (FDR < 3.1 x 10^-5^) that are putatively regulated by Rad53. This identified several expected terms –related to DNA replication, chromatin organization, and checkpoint control – as highly significantly enriched (Fig 1H). Unexpectedly, the term ‘*positive regulation of metabolic process’* was enriched among the top categories with the false discovery rate (FDR) = 3.17 x 10^-6^ (Fig 1H). While all other terms were associated with cell cycle progression, only this category reflected cellular metabolic capacity, and raised the possibility that Rad53 might be associated with metabolic regulation — a function that has not been previously explored (Fig 1I).

### *rad53Δ* cells exhibit altered glycolysis and increased TCA flux

We therefore examined the overall metabolic state of *rad53Δ* cells in standard high glucose medium, by quantitatively assessing central carbon metabolism intermediates – from glycolysis, the pentose phosphate pathway and tricarboxylic acid (TCA) cycle – in *rad53Δ* and WT cells (see methods), using highly targeted, quantitative LC/MS/MS approaches (Fig 2A)^29^. Central carbon metabolism collectively reflects nutrient availability, bioenergetic demand, and biosynthetic flux^30^, and changes in core metabolites sensitively indicate changes in cellular metabolic state. These analyses revealed increased amounts of glycolytic and Pentose phosphate pathway (PPP) intermediates (Fig 2B, S2A), as well as significantly increased amounts of the tricarboxylic acid (TCA) cycle intermediates in the *rad53Δ* cells compared to WT cells growing in the same conditions (Fig 2B, S2A). This increase in TCA cycle intermediates was particularly notable, since *S. cerevisiae* cells growing in high glucose are Crabtree positive and highly glycolytic, and typically repress their mitochondrial outputs^31^. This significant increase in mitochondrial TCA intermediates observed in *rad53Δ* cells was therefore unexpected. To more systematically address this finding, and distinguish between increased metabolite accumulation due to reduced utilization, vs higher production through the TCA, we next estimated glucose flux into the TCA cycle, using stable-isotope (U^13^C_6_ glucose) based carbon isotope tracing^32,33^. For this, we spiked cells growing log phase with 1% U^13^C_6_ glucose for 10 min, and estimated the relative fraction of stable-isotope incorporation into TCA intermediates (Fig 2A, more details in Fig S2B). Notably, we observed a substantial increase in the relative carbon label incorporation from glucose derived pyruvate into the indicated TCA metabolites in *rad53Δ* cells (Fig 2C).

**Fig 2.**
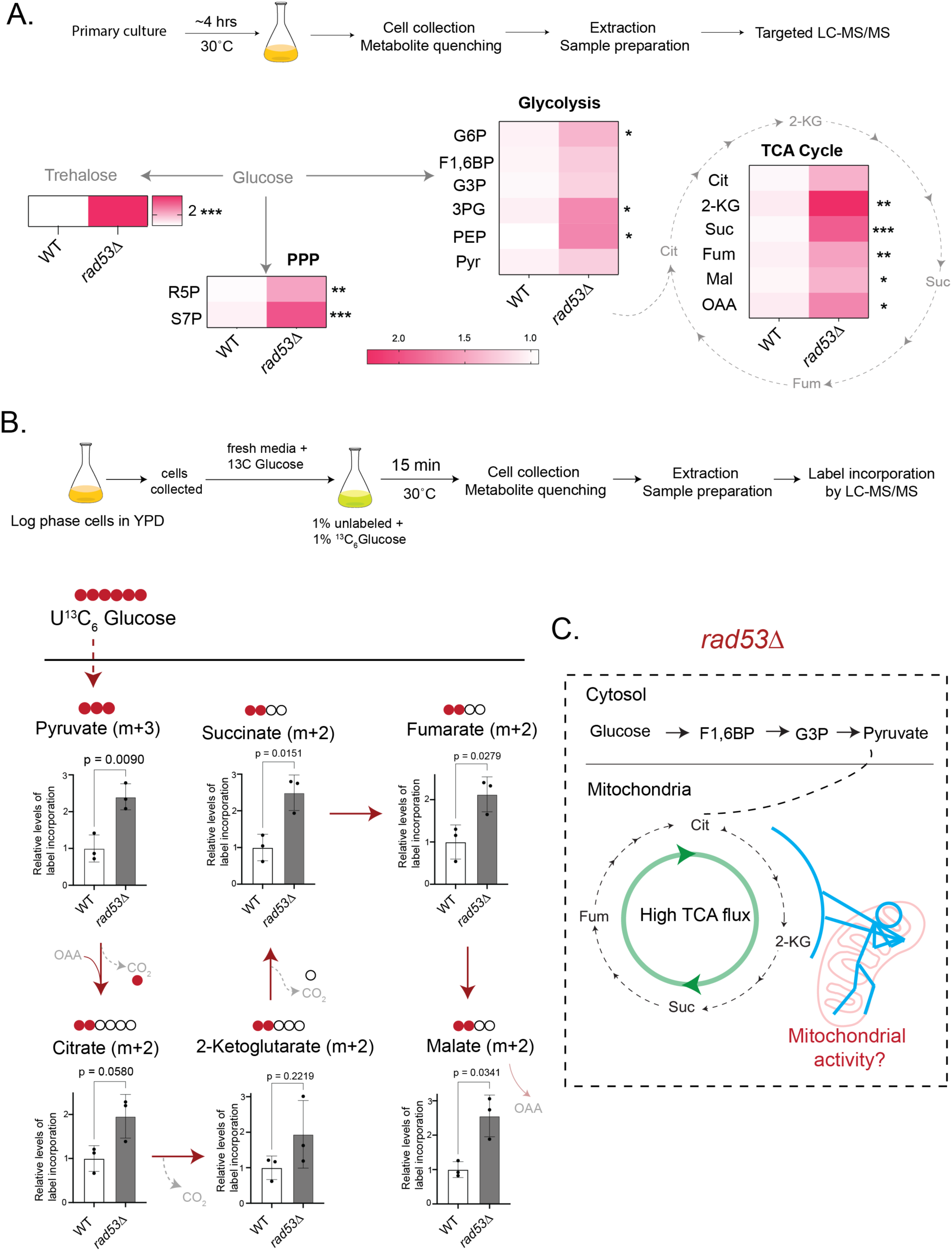
*rad53Δ* cells exhibit rewired metabolism with increased flux and allocations towards the TCA cycle. **A.** Quantitative LC–MS/MS-based assessment of central carbon metabolism in WT and *rad53Δ* cells. A schematic of the experimental workflow is shown alongside a heat map depicting the relative abundance of metabolites from glycolysis, the tricarboxylic acid (TCA) cycle, and the pentose phosphate pathway (PPP). Heat map represents mean values for n ≥ 3, *p<0.05, **p<0.01, ***p<0.001, ****p<0.0001, two-tailed, unpaired Welch’s t-test. Also, see Fig S2A for quantified bar graphs of individual metabolite data. **B.** Stable isotope tracing using ^13^C_6_ glucose to assess carbon flux into TCA cycle metabolites in WT and *rad53Δ* cells. A schematic of the ^13^C_6_ glucose labelling and metabolite analysis workflow is shown, followed by quantification of ^13^C incorporation into TCA cycle metabolites. *rad53Δ* cells exhibit significantly increased carbon flux through the TCA cycle compared with WT cells. Data are displayed as means ± SD, n = 3. *p<0.05, **p<0.01, ***p<0.001, ****p<0.0001, two-tailed, unpaired Welch’s T test. Also, se Fig S2B for carbon tracing information. **D.** Working hypothesis to determine the metabolic state of *rad53Δ* cells, where altered mitochondrial function exerts a metabolic ‘pull’ on glucose-derived carbon, towards increased flux into the TCA cycle.

These data establish that *rad53Δ* cells have a significant increase in their TCA cycle flux, coming from glucose derived pyruvate entering into the mitochondrial TCA cycle. These data therefore reveal an unexpected metabolic rewiring in *rad53Δ* cells growing in high glucose, with increased pyruvate allocations towards the mitochondrial TCA cycle, and high TCA flux. This high mitochondrial TCA flux in *rad53Δ* cells suggest a ‘pull’ towards increased mitochondrial metabolism.

### Quantitative proteomics reveal a remodeled mitochondrial proteome in *rad53Δ* cells

We therefore sought to examine mitochondrial function and composition in *rad53Δ* cells. As a preliminary, qualitative readout for mitochondrial inner membrane amounts, we first used NAO (nonyl acridine orange), a strong, fluorescent cardiolipin-binding dye^34^. The NAO fluorescence in *rad53Δ* was significantly higher than those of WT cells (Fig 3A and S3A), suggesting an increase in total mitochondrial cardiolipin amount. We therefore next developed an unbiased, data-independent acquisition (DIA) based quantitative proteomic approach, in order to compare the mitochondrial proteome between *rad53Δ* and wild-type cells (Fig 3B). To quantitatively investigate the changes in mitochondrial composition, and reduce background from cytoplasmic proteins and organelles, we first enriched the total mitochondrial fractions in WT and *rad53Δ* cells, by using extensive mito-fractionation protocols established earlier^31,35,36^ (Fig 3B). Equal amounts of purified total mitochondrial protein (100 µg) from each isolated biological replicate were comparatively analyzed for this experiment, using three replicates for WT and *rad53Δ* mitochondrial isolates respectively (Fig S3B). The resulting mitochondrial fractions were resolved on SDS-PAGE gels, followed by in-gel trypsin digestion and sample analysis on Orbitrap Fusion Tribid mass spectrometer (Thermo Fisher Scientific). Data-independent acquisition (DIA)-based proteomics was used for protein quantification, with identification and quantification performed using DIA-NN^37^, allowing relative-quantitative comparisons of identified mitochondrial protein abundances between wild-type and *rad53Δ* cells (Fig 3B). Through this, we were able to quantitatively compare changes in the respective mitochondrial proteomes, for the ∼same amount of input mitochondrial protein (Fig 3B).

**Fig 3.**
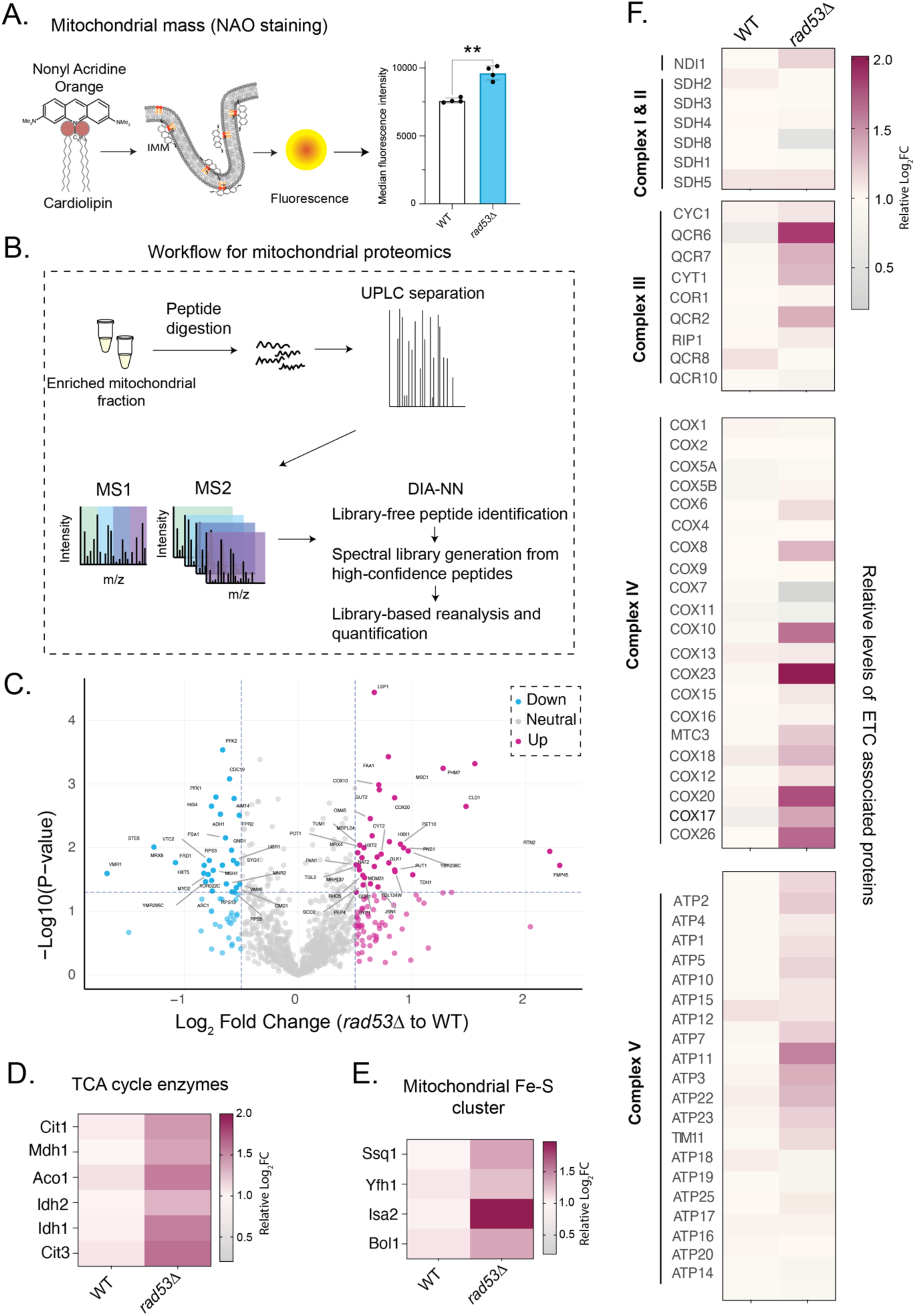
Altered mitochondrial proteome and ETC composition in *rad53Δ* cells. **A.** (Left) Schematic illustrating the binding of NAO to cardiolipin in the inner mitochondrial membrane, and resulting fluorescence increase based on cardiolipin abundance, as a method to quantify mitochondrial amounts. (Right) Relative median 10-N-nonyl acridine orange (NAO) fluorescence intensity in WT and *rad53Δ* cells. Data displayed as means ± SD, n = 4. **p<0.01, two-tailed, unpaired Welch’s T test. Also see Fig S3A for flow cytometry-based intensity distribution for NAO fluorescence, to reflect differences in mitochondrial content. **B.** Workflow for mitochondrial proteomics. Enriched mitochondrial fractions were subjected to peptide digestion, followed by UPLC separation and data-independent acquisition (DIA) mass spectrometry. Peptides were identified and quantified using DIA-NN. For all workflows, three independent biological replicates were used for each background. Also see Fig S3B, to show input amounts of total isolated mitochondria, across all the independent replicates. **C.** A volcano plot depicting differences in protein abundance in *rad53Δ* cells relative to WT cells. Log₂ fold changes were calculated from the mean protein abundances across three independent biological replicates per condition. P values were calculated using a two-tailed, unpaired Welch’s T-test. Proteins with a log₂ fold change > 0.5 or < −0.5 are highlighted (n = 3 independent biological replicates per condition). Also see Fig S3C and S3D for GO analysis of biological processes of differentially abundant proteins. **D.** Heat map depicting relative abundances of enzymes of the TCA cycle proteins in *rad53Δ* and WT cells. Heat map showing the median values across three independent biological replicates (n = 3). **E.** Heat map depicting relative abundances of the Fe-S cluster proteins in *rad53Δ* and WT cells. Heat map showing the median values across three independent biological replicates (n = 3). **F.** Heat map showing relative abundances of the Electron transport chain (ETC) associated proteins in *rad53Δ* and WT cells. Heat map showing the median values across three independent biological replicates (n = 3). Also see Fig S3E, S3F for bar graphs showing the quantification of selected individual proteins.

For subsequent data analysis, we filtered the proteomics dataset to retain only mitochondrial proteins, based on annotations from the Saccharomyces Genome Database (SGD), and curated yeast mitochondrial proteome datasets^38–40^ (also see Methods for more details). Following this and an extensive manual curation, the analyses yielded a final set of 1,073 mitochondrial proteins quantitatively assessed after our enrichment-based purification (Table S3). To identify abundance changes in *rad53Δ* relative to WT cells, we applied a fold-change cutoff of ±0.5 Log₂FC (i.e. >1.4-fold change, across replicates), revealing 150 mitochondrial proteins that were significantly increased or decreased in *rad53Δ* cells (Fig 3C, Table S3). Interestingly, while several mitochondrial proteins were substantially increased in *rad53Δ* cells, several others were significantly reduced, suggesting non-uniform changes in the overall mitochondrial proteome of *rad53Δ* cells. Gene ontology (GO) enrichment (ShinyGO) analysis of the differentially abundant proteins indicated clear functional trends in the *rad53Δ* cell mitochondrial proteome. Proteins increased in *rad53Δ* cells were enriched for specific mitochondrial and metabolic processes, such as short-chain fatty acid metabolism, mitochondrial respiratory chain complex assembly, and broader carboxylic metabolic pathways, as well as mitochondrial organization (Fig S3C, Table S4). In contrast, the downregulated proteins were predominantly associated with ATP generation from ADP and nucleotide phosphorylation processes (Fig S3D, Table S4).

To more meaningfully resolve pathway-level changes within the mitochondrial proteome, we manually re-grouped mitochondrial proteins into major functional categories - the electron transport chain (ETC), TCA cycle, Fe–S (Iron-Sulfur) cluster biogenesis, mitochondrial ribosomes, and the fission-fusion machinery. Proteins belonging to these categories were extracted based on SGD annotations, and their relative abundances in *rad53Δ* vs WT was compared. This analysis revealed a pronounced overall increase in relative amounts of proteins involved in the TCA cycle, the iron-sulfur clusters (Fig 3D, Fig 3E), along with an increase in cardiolipin remodeling protein CLD1 (cardiolipin-specific deacylase 1) in *rad53Δ* cells (Fig 3D, 3E; S3E; Table S5).

We next examined the relative abundance of the mitochondrial electron transport chain (ETC)-associated proteins across the respiratory complexes I-V. Notably, our analyses revealed that these proteins displayed a non-uniform pattern of increase – several ETC subunit proteins were significantly increased in *rad53Δ* cells, but many remained unchanged, and a subset were reduced in *rad53Δ* relative to wild type (Fig 3F, S3F; Table S5). Among the increased proteins, several annotated components associated with complex IV and V showed significant increase, however this was not uniform across all the complex sub-units. (Fig 3F, S3F). Collectively, instead of an increase across all the ETC complex proteins in *rad53Δ* cells, some ETC components variably increased, while others were unchanged or even reduced. In contrast, the mitochondrial ribosomal subunits, or the components of the mitochondrial fission and fusion machinery, did not show any statistically significant changes, and a modest upward trend (Fig S3G-I; Table S5).

Overall, these data reveal that *rad53Δ* cells exhibit an increase in select mitochondrial metabolic and membrane-associated pathways, with a particular enrichment of TCA cycle, and cardiolipin-related proteins, alongside a reorganization of the ETC subunits with non-uniform increases in their amounts.

### *rad53Δ* cells have high mitochondrial potential with unaltered respiration, high ROS and oxidative stress

Mitochondrial outputs rely on coordinated electron transfer through the ETC and the formation of a proton gradient across the mitochondrial membrane, resulting in the terminal electron acceptance by oxygen (oxygen consumption) at complex IV, coupled with the proton motive force driving the ATP synthase complex (Complex V). Since *rad53Δ* cells exhibited non-uniform changes in the abundance of ETC sub-units (Fig 3), we next asked whether these different functional steps of mitochondrial activity were affected in *rad53Δ* relative to WT cells. We measured multiple mitochondrial outputs independently - the amounts of a key ETC component (cytochrome c oxidase in Complex IV)^41^, inner mitochondrial membrane potential, oxygen consumption, and cellular ATP production, in WT and *rad53Δ* cells. We first compared the relative levels of Cox2 protein (cytochrome c oxidase). We observed that the amounts of Cox2 in *rad53Δ* cells was substantially higher than in WT cells (Fig 4A). Separately, we observed increased levels of Porin (mitochondrial outer membrane porin, Por1) in *rad53Δ* cells (Fig S4A). Next, we assessed mitochondrial membrane potential using a potentiometric dye, Mitotracker Red (CMXRos). *rad53Δ* cells exhibited significantly higher mitochondrial potential compared to wild-type cells (Fig 4B). As a control, cells were treated with sodium azide, an ETC complex IV inhibitor, which lead to marked reduction in fluorescence, confirming that the measured signal was specific to mitochondrial potential (Fig S4B). Together, these data indicate significantly higher mitochondrial potential and activity of the electron transport chain in *rad53Δ* cells (Fig 4A, 4B).

**Fig 4.**
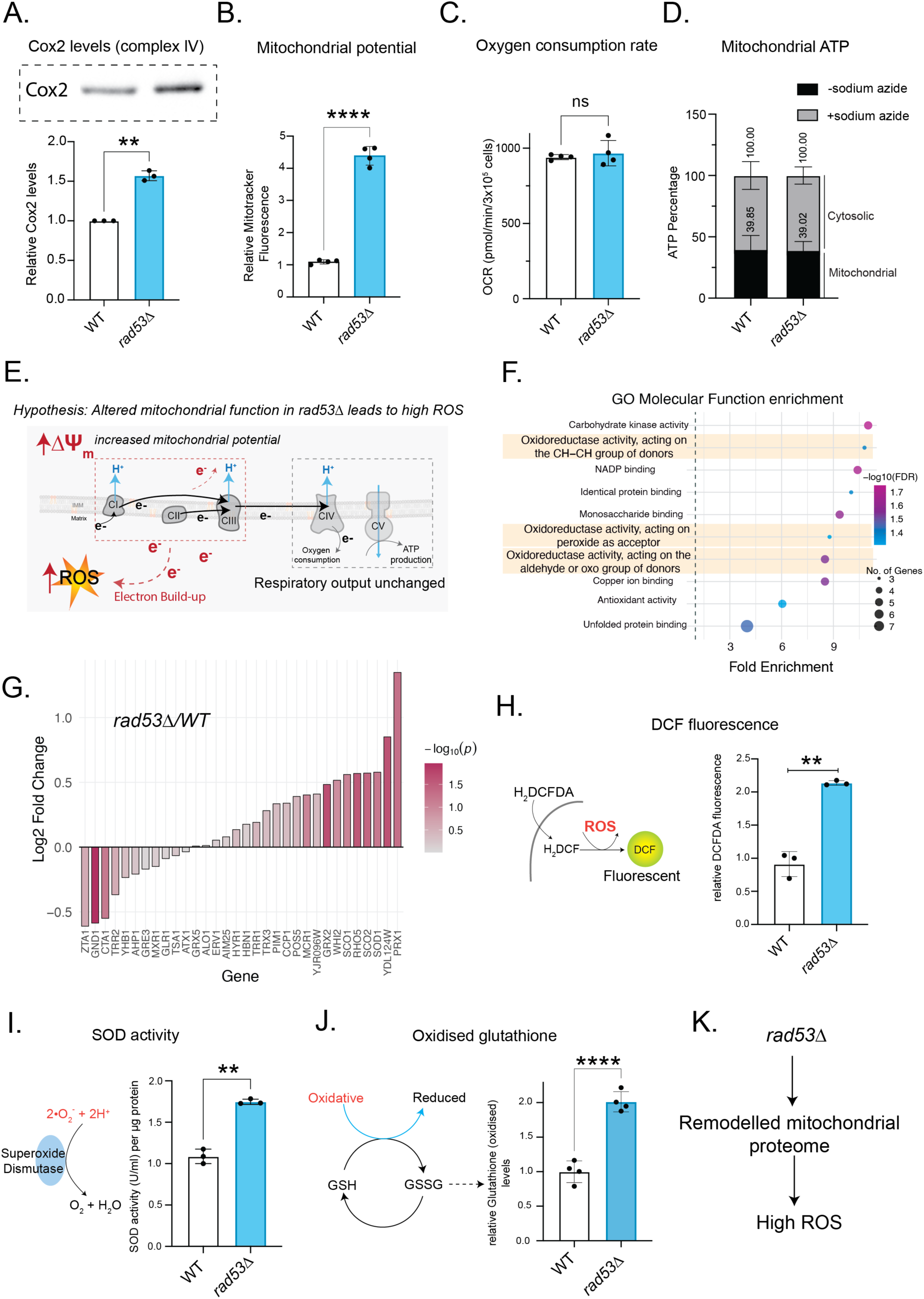
Altered mitochondrial function without increased oxygen consumption resulting in high ROS in *rad53Δ* cells. **A.** Relative amounts of total cytochrome c oxidase subunit 2 (Cox2) in WT and *rad53Δ* cells determined by Western blotting. Representative immunoblot and quantification of Cox2 levels are shown. Data are displayed as means ± SD, n = 3. **p<0.01, two-tailed, unpaired Welch’s T test. Also see Fig S4A for levels of the outer mitochondrial membrane Por1 protein. **B.** Relative mitochondrial membrane potential in WT and *rad53Δ* cells measured using the potentiometric dye MitoTracker Red CMXRos. *rad53Δ* cells exhibit significantly higher mitochondrial potential than WT cells. Data are displayed as means ± SD, n = 4. ****p<0.0001, two-tailed, unpaired Welch’s T test. Also see Fig S4B for relative mitochondrial membrane potential with or without inhibition of complex IV with sodium azide. **C.** Relative basal oxygen consumption rate (OCR) in WT vs *rad53Δ* cells, quantified from Seahorse extracellular flux analyzer. Data are displayed as means ± SD, n = 4. ns denotes non-significant difference, two-tailed, unpaired Welch’s T test. Also see Fig S4C for OCR over time. **D.** Relative contribution of mitochondrial ATP to total cellular ATP in WT and *rad53Δ* cells, measured in the presence or absence of sodium azide. Data are presented as percentages of total cellular ATP, n = 3. Also see Fig S4D for total ATP concentrations, and the total cytosolic ATP concentrations. Fig S4E shows ATP data in cells treated with oligomycin. **E.** A hypothetical model illustrating altered mitochondrial electron transport in *rad53Δ* cells. Enhanced electron flow through the upstream electron transport chain complexes may result in increased electron leakage and ROS generation, while downstream respiratory flux, oxygen consumption, and ATP production would remain unchanged. **F.** Gene Ontology (GO) molecular function enrichment analysis of differentially expressed mitochondrial proteins in *rad53Δ* cells. Enriched terms are ranked by fold enrichment, with oxidoreductase activity related terms among the most highly enriched functions. Significance is indicated by −log₁₀(FDR). **G.** Relative abundance of mitochondrial proteins associated with oxidative stress responses in *rad53Δ* vs WT cells. Log₂ fold changes (*rad53Δ* relative to WT) were derived from the quantitative proteomic dataset for n = 3. P values are calculated by two-tailed, unpaired Welch’s T test. **H.** ROS levels in WT and *rad53Δ* cells were quantified using the ROS-sensitive fluorescent dye H₂DCFDA. The schematic on the left illustrates the mechanism of ROS-dependent oxidation of H₂DCFDA leading to the fluorescent product DCF. Data displayed as means ± SD, n = 3. **p<0.01, two-tailed, unpaired Welch’s T test. **I.** SOD activity in WT and *rad53Δ* cells was quantified to assess the cellular response to elevated oxidative stress. The schematic on the left illustrates the dismutation of superoxide radicals by superoxide dismutase (SOD), generating hydrogen peroxide and molecular oxygen. Data displayed as means ± SD, n = 3. **p<0.01, two-tailed, unpaired Welch’s T test. **J.** Relative amounts of redox responsive oxidized glutathione in WT vs *rad53Δ* cells., Data displayed as means ± SD, n = 4. ****p<0.0001, two-tailed, unpaired Welch’s T test. Also see Fig S4F for reduced glutathione amounts. **K.** Loss of Rad53 is associated with mitochondrial ETC imbalance and elevated cellular ROS.

We next assessed the basal oxygen consumption rate (OCR) using a Seahorse flux analyzer, in WT and *rad53Δ* cells. Surprisingly, the *rad53Δ* cells did not show any significant increase in OCR compared to WT cells (Fig 4C, S4C), indicating that the mitochondria in *rad53Δ* did not have increased oxygen consumption. Since oxygen consumption by mitochondria is coupled to ATP generation by the ATP synthase, we next assessed the relative ATP production at complex V, and compared the total and relative mitochondrial-contribution towards ATP amounts in WT and *rad53Δ* cells respectively. For this, we estimated total ATP in these cells, with and without a Complex IV inhibitor (sodium azide), to quantify relative mitochondrial contributions towards total ATP, as described earlier ^31,42^. Notably, both total and mitochondrial ATP levels were comparable between WT and *rad53Δ* cells, with no significant differences observed (Fig 4D, S4D). These data are also consistent with the unchanged OCR observed earlier (Fig 4C) in *rad53Δ* cells. Finally, we hypothesized that it may be possible for the ATP synthase (complex V) itself to hydrolyze ATP and generate mitochondrial membrane potential. To determine whether this factor contributed to the increased membrane potential in *rad53Δ*, we treated these cells with oligomycin (an ATP synthase inhibitor). Inhibiting ATP synthase led to a further increase in mitochondrial membrane potential in both wild-type and mutant cells to a similar extent, indicating that the increased mitochondrial potential in *rad53Δ* is not likely to be driven by ATP hydrolysis (Fig S4E). Therefore, these data collectively revealed that *rad53Δ* cells exhibit unusual mitochondrial functioning – having a non-uniform increase in select ETC subunits, and substantially increased mitochondrial potential, but no increase in the final mitochondrial ETC outputs – oxygen consumption and ATP synthesis. These are consistent with an altered mitochondrial proteome observed in *rad53Δ*.

One possible consequence of such an imbalanced mitochondrial proteome in *rad53Δ*, without increased oxygen consumption or ATP generation, would be an increased possibility of electrons leaking, leading to the generation of reactive oxygen species (ROS) as is well established^42,43^, and illustrated in Fig 4E. If this were so, a consequence of these altered mitochondria in *rad53Δ* cells would be a substantial increase in ROS and subsequent oxidative stress.

We therefore re-inspected our proteome data, to assess if the mitochondrial proteome of *rad53Δ* cells exhibited signatures associated with oxidative stress due to such changes in mitochondrial function. Gene Ontology (GO) molecular function analysis revealed strong enrichment of oxidoreductases and oxidation-related proteins among the differentially expressed mitochondrial proteins in *rad53Δ* cells. (Fig 4F). We next manually curated mitochondrial proteins that are critical in managing an oxidative stress response in cells (annotated in SGD) (Table S6), and assessed their fold-changes from our proteome data (Fig 4E). Notably, several oxidative stress related proteins, including peroxidases and superoxide dismutases, are significantly increased in *rad53Δ* cells (Fig 4G). Collectively, these data suggested that *rad53Δ* cells exhibited features of encountering high ROS.

To functionally assess this, we first estimated the relative cellular ROS amounts of WT and *rad53Δ* cells. For this, we used DCFDA, a cell-permeable dye that is oxidized (by ROS) and cleaved by esterases to increase its fluorescence, and which reliably indicates increased intracellular ROS^42^. *rad53Δ* cells showed significantly increased DCFDA fluorescence compared to WT cells, indicating very high ROS in these cells (Fig 4H). Next, we measured multiple biochemical outputs that would be consistent with cells responding to increased reactive oxygen species. We first assessed superoxide dismutase (SOD) activity, since SOD is required to remove ROS. *rad53Δ* cells show significantly increased SOD activity compared to WT cells (Fig 4I), consistent with increased DCFDA fluorescence. Next, we estimated the amounts of glutathione, a bulk reductant that increases in cells as a response to high oxidative stress. Total as well as oxidized glutathione levels substantially increased in *rad53Δ*, consistent with these cells having higher ROS (Fig 4J; S4F). Collectively, our data reveal that *rad53Δ* cells have altered mitochondrial proteomes and high membrane potential without increased respiration, resulting in high ROS and oxidative stress in these cells (Fig 4K).

### High mitochondrial potential and oxidative stress in *rad53Δ* are kinase-dependent and distinct from the DNA-damage Rad53/Sml1 axis

We next asked whether the observed mitochondrial and oxidative stress phenotypes associated with the complete loss of the Rad53 protein depend on its kinase catalytic function. The Rad53 kinase domain extends from amino acid residue 198 to 466, with a critical catalytic region that includes K227. We therefore used genome-editing, focusing on the region that contained the critical catalytic K227 residue within the kinase domain. We engineered a catalytically-dead strain where a stretch of amino acids in the catalytic region alone were mutated, with the final sequence as shown in Fig 5A. This strain is referred to as the kinase-deficient Rad53 mutant (Rad53*^KD^*) for the remainder of this manuscript. We confirmed that this strain was hypersensitive to DNA-damage inducing agents (MMS) and replication stress (hydroxyurea), consistent with impaired Rad53 catalytic function (Fig S5A, S5B). We next assessed whether Rad53*^KD^*was sufficient to reproduce the phenotypes observed in *rad53Δ* cells. The kinase-deficient mutant Rad53*^KD^* showed increased mitochondrial membraned potential (as measured using CMX-ROS Mitotracker red) (Fig 5B), and increased oxidative stress (Fig 5C), and exhibited a slower growth (Fig 5D), closely phenocopying the *rad53Δ* cells. Together these findings suggest that the mitochondrial and oxidative stress phenotypes associated with Rad53 loss depend on the Rad53 kinase catalytic function.

**Fig 5.**
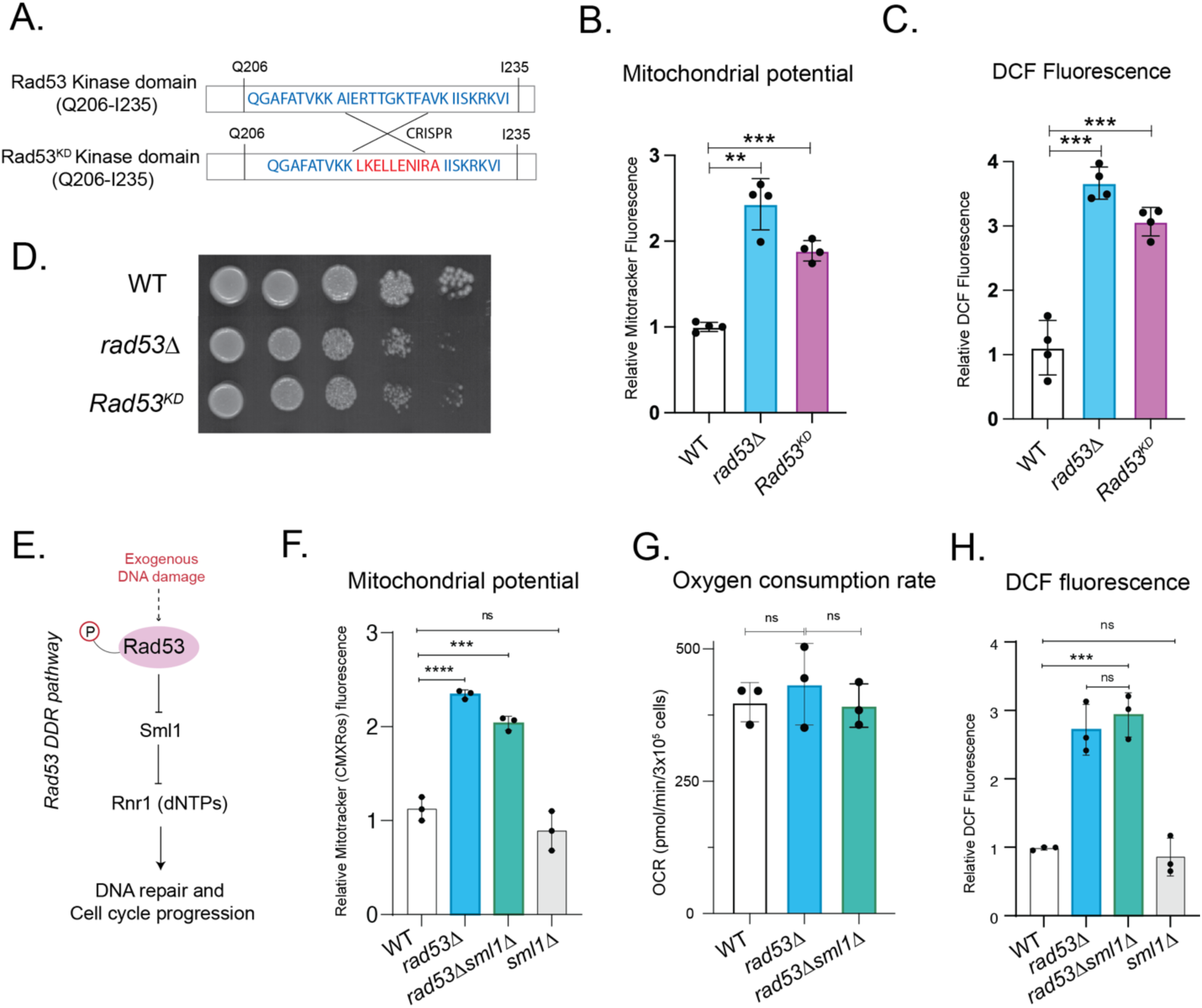
Altered mitochondrial outputs and high ROS in *rad53Δ* depends on kinase function and is distinct from the Rad53/Sml1/DNA damage axis: **A.** Illustration of the amino acid sequence of the Rad53 kinase domain region in WT cells, and the CRISPR-generated Rad53^KD^ mutant, with the altered amino acid sequence in the mutant shown in red. **B.** Relative mitochondrial membrane potential in WT, *rad53Δ*, and Rad53^KD^ cells, measured using MitoTracker Red CMXRos fluorescence. Data displayed as means ± SD, n = 4, **p<0.01, ****p<0.0001, two-tailed, unpaired Welch’s T test. **C.** Intracellular ROS levels in WT, *rad53Δ*, and Rad53^KD^ cells, quantified using the ROS-sensitive fluorescent dye DCFDA. Data displayed as means ± SD, n = 4, independent replicates. ***p<0.001, two-tailed, unpaired Welch’s T test. **D.** Growth of WT, *rad53Δ* and Rad53^KD^ cells in YPD medium on solid agar. The *rad53Δ* and Rad53^KD^ cells exhibited reduced growth, compared to WT. Also see Fig S5A and S5B for growth in the presence of DNA damaging agents. **E.** Schematic illustrating the well-established Rad53 DDR pathway where Rad53 is activated upon DNA damage, and inhibits Sml1, which is an inhibitor of the ribonucleotide reductase Rnr1, to collectively relieve Rnr1 inhibition. Also see Fig S5C and Fig S5D for the generation and genotyping of the *rad53Δ/ sml1Δ* strains. **F.** Mitochondrial membrane potential in *rad53Δ* and *rad53Δ/sml1Δ* cells, compared to WT or *sml1Δ* cells, as measured using MitoTracker Red CMXRos. Data displayed as means ± SD, n = 3, **p<0.01, ****p<0.0001, ns denotes non-significant difference, two-tailed, unpaired Welch’s T test. **G.** Basal oxygen consumption in *rad53Δ* and *rad53Δ/sml1Δ* cells compared to WT cells. Also see Fig. S5E for oxygen consumption over time. Data displayed as means ± SD, n = 3, ns denotes non-significant difference, two-tailed, unpaired Welch’s T test. **H.** Relative DCFDA fluorescence in *rad53Δ* and *rad53Δ/sml1Δ* cells compared to WT and *sml1Δ* cells. Data displayed as means ± SD, n = 3, ***p<0.001, ns denotes non-significant difference, two-tailed, unpaired Welch’s T test.

We therefore next asked whether the Rad53 dependent restraint on mitochondrial outputs and ROS occurred through the canonical DNA-damage Rad53-Sml1 axis. To recall, during DNA damage, hyperphosphorylated Rad53 phosphorylates and represses Sml1, which in turn results in the activation of Rnr1 and DNA repair (Fig 5E). To address this, we generated *SML1* knock-out strains in WT (*sml1Δ*) and *rad53Δ* (*rad53Δ/sml1Δ*) cells. For this, we generated diploid *sml1Δ* strains, in which a single copy of *RAD53* was knocked-out, and haploid *rad53Δ/sml1Δ* strains were subsequently obtained via sporulation and dissection of the daughter spores (Fig S5C). Spores were confirmed for the *rad53Δ/sml1Δ* genotype by PCR (Fig S5D).

Using the *rad53Δ*/*sml1Δ* and *sml1Δ* strains, we next compared what the mitochondrial state and ROS levels were in these respective strains. For this, we first assessed the relative mitochondrial membrane potential of *rad53Δ*/*sml1Δ* cells (with methods as described earlier). Notably, the *rad53Δ and rad53Δ*/*sml1Δ* cells were largely indistinguishable, showing comparable, high mitochondrial potential, (Fig 5F). In contrast, the WT and *sml1Δ* cells had lower (comparable) potential (Fig 5F), indicating that the *sml1Δ* cells resembled WT and not *rad53Δ/sml1Δ* cells. Next, we measured oxygen consumption rate (OCR) of the *rad53Δ*/*sml1Δ* cells. We observed that *rad53Δ*/*sml1Δ* cells had similar oxygen consumption as *rad53Δ* cells (Fig 5G; S5E), while the *sml1Δ* cells resembled WT cells. Finally, we finally compared ROS levels in *rad53Δ/sml1Δ* cells with *rad53Δ* and WT cells. We assessed the relative amount of ROS in *rad53Δ*/*sml1Δ* cells, using DCFDA fluorescence. Consistent with the mitochondrial outputs, the *rad53Δ*/*sml1Δ* cells show increased ROS levels, closely resembling *rad53Δ* cells (Fig 5H). In contrast, *sml1Δ* cells resembled WT cells, with comparable, lower ROS (Fig 5H). Finally, we compared the growth of the *rad53Δ/sml1Δ* cells, along with the *sml1Δ* cells. The *rad53Δ/sml1Δ* cells showed growth that phenocopied *rad53Δ* cells, while in contrast *sml1Δ* resembled WT cells (Fig S5F).

Collectively, these findings indicate that the role of Rad53 in restraining mitochondrial outputs and ROS are distinct from the Rad53/Sml1 DNA damage response axis. Thus, Rad53 kinase activity is critical to maintain mitochondrial balance and limit oxidative stress, and this function is distinct from its canonical role in the DNA damage response through Sml1 regulation.

### Scavenging ROS in *rad53Δ* is sufficient to rescue and restore G2/M progression

Finally, we asked if the observed G2/M and cell cycle progression delay, cell size heterogeneity, and slow growth (as established in Fig 1) was caused by the high ROS and oxidative stress in the *rad53Δ* cells. For this, we first assessed how resistant WT vs *rad53Δ* cells were to exogenous oxidative stress. When grown in the presence of hydrogen peroxide, *rad53Δ* cells were acutely hypersensitive to hydrogen peroxide (Fig 6A), showing almost no growth, in contrast to WT cells which were largely unaffected for growth at these concentrations of hydrogen peroxide. We next assessed if the delayed G2/M progression in in *rad53Δ* cells was caused by the high ROS, coming from the imbalanced mitochondrial proteome and outputs. To assess this, we treated cells with a potent anti-oxidant, ascorbic acid, which directly scavenges electrons to reduce ROS and resultant oxidative stress^42,44^. We first measured the relative ROS levels in *rad53Δ* cells upon adding ascorbic acid. We observed that the ROS levels in *rad53Δ* cells are substantially reduced upon ascorbic acid addition (Fig 6B), to levels comparable to WT cells. Next, we extensively assessed budding time and cell division in *rad53Δ* cells, with or without ascorbic acid addition. The mean budding time in *rad53Δ* cells decreased from 149 minutes to 98 minutes in the presence of ascorbic acid (Movie S5), and the median budding time of *rad53Δ* cells reduces from 140 minutes to ∼70 minutes, largely comparable to WT cells growing in YPD. This revealed a ∼near complete rescue in the budding time of *rad53Δ* cells treated with the anti-oxidant (Fig 6C; S6A, Movie S3, S4, S5). A small number of *rad53Δ* cells remained permanently arrested and incapable of resuming division (Fig S6B, Movie S6), which we infer is likely due to accumulated, irrecoverable damage due to high ROS. Since we had earlier observed that a significant subpopulation of *rad53Δ* (∼6-7%) were substantially enlarged (Fig 1F), we next asked whether the total fraction of highly enlarged-cells changed upon ascorbic acid treatment. Notably, ascorbic acid treatment significantly reduced the proportion of enlarged *rad53Δ* cells, and the fraction of cells with projected areas >30 µm² decreased more than two-fold (Fig 6D, S6C). Therefore, scavenging the produced ROS using antioxidant treatment reduced the accumulation of abnormally large *rad53Δ* cells in the population. Finally, we assessed overall cell growth (based on biomass increase) after ascorbic acid treatment. Ascorbic acid addition substantially reduced the lag phase and restored the rate of biomass increase in *rad53*Δ cells (Fig 6E, S6D).

**Fig 6.**
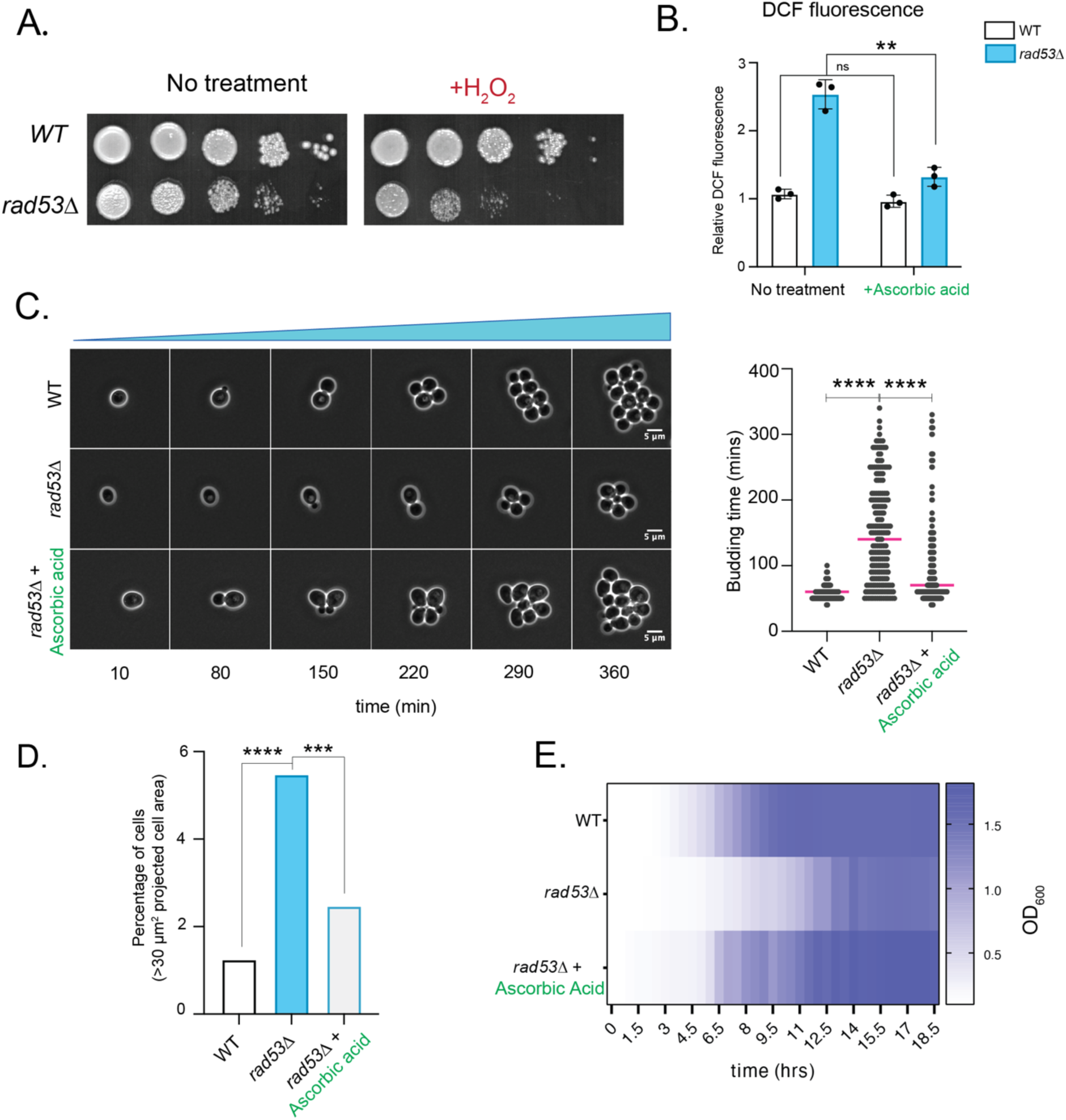
Anti-oxidant treatment to reduce ROS in *rad53Δ* cells rescues G2/M and cell cycle progression. **A.** Growth of WT and *rad53Δ* cells in the presence of hydrogen peroxide (H₂O₂). *rad53Δ* cells exhibit hyper-sensitivity to oxidative stress compared to WT cells. **B.** Intracellular ROS levels in WT and *rad53Δ* cells following treatment with the antioxidant ascorbic acid, quantified using the ROS-sensitive fluorescent dye DCFDA. Ascorbic acid treatment reduced ROS levels in *rad53Δ* cells. Data displayed as means ± SD, n = 3, **p<0.01, ns denotes non-significant difference, two-tailed, unpaired Welch’s T test. **C.** (Left) Representative time-lapse images of WT and *rad53*Δ cells grown in the presence or absence of the antioxidant ascorbic acid, illustrating the restoration of budding kinetics following ROS reduction. Time (min) is indicated below each frame. Scale bar, 5 μm. (Right) Mean and median budding times of WT and *rad53Δ* cells in the presence or absence of ascorbic acid, determined by time-lapse microscopy. Ascorbic acid treatment reduced the mean budding time of *rad53Δ* cells from 149 min to 98 min. Individual data points are shown, with the median indicated. ****p<0.0001, two-tailed Mann–Whitney U test. WT, n = 140; *rad53*Δ, n = 180; *rad53*Δ + ascorbic acid, n = 185. Also see Fig S6A for interquartile range (IQR) of budding time and budding time distribution. Also see Movie S3-S5 for real-time data of cell division. **D.** Proportion of enlarged cells (projected area >30 μm²) in WT, *rad53Δ*, and ascorbic acid-treated *rad53Δ* cells. Antioxidant treatment markedly reduced the enlarged-cell subpopulation in *rad53Δ* cells. Statistical significance was determined using Fisher’s exact test (****p< 0.0001; exact *P* values indicated; WT, *n* = 800; *rad53Δ, n* = 900*; rad53Δ +* ascorbic acid, *n* = 1000, pooled from three independent paired experiments). Also see Fig S6C for cell size projections. **E.** Growth of WT and *rad53Δ* cells in liquid culture in the presence of the ROS scavenger ascorbic acid. The heat map represents mean optical density (OD) at each time point and shows the rescue of the delayed growth of *rad53Δ* cells after ascorbic acid treatment. Data represent means from three biological replicates (n = 3). Also see Fig S6E for continuous growth curve data for the respective conditions.

Collectively, scavenging ROS and restoring oxidative balance in *rad53*Δ cells rescued cell division, cell cycle progression and cell growth largely comparable to WT cells.

## Discussion

In this study, we discover that the checkpoint kinase Rad53/Chk2 enforces a metabolic checkpoint for cell cycle progression. In cells growing in high glucose, Rad53 restrains mitochondrial outputs to limit ROS production, allowing timely G2/M progression (Fig 6F). This role in G2/M progression is distinct from the well-studied Rad53/Sml1 DNA damage axis, which is activated in response to acute DNA damage by exogenous damaging agents. Our findings substantially broaden the role of Rad53 kinase beyond its established function in the DNA damage response, and reveal a role in coordinating mitochondrial activity to restrain ROS induced oxidative stress that can impede growth. This allows normal G2/M cell cycle progression and rapid growth, as illustrated in the model in Fig 7.

**Fig 7.**
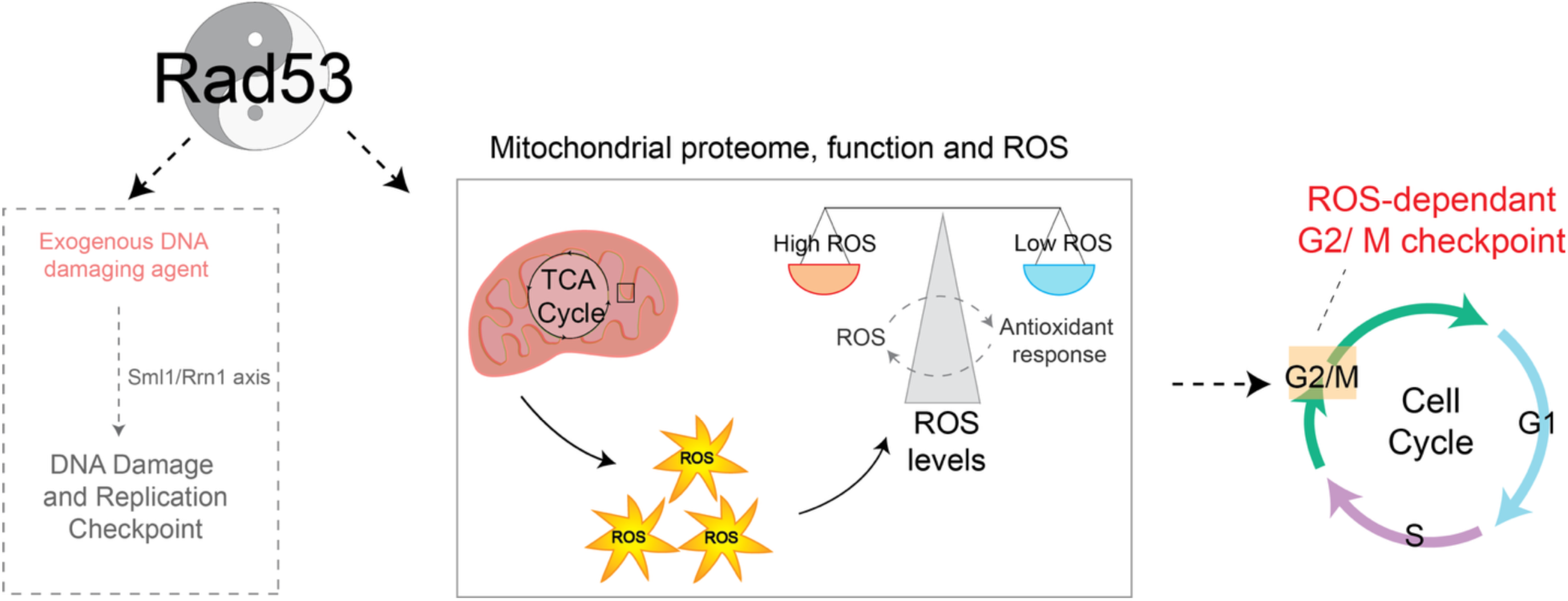
Rad53 restrains mitochondrial function and ROS to impose a ROS-dependent G2/M checkpoint. Model illustrating: (Left) Rad53 functions as a DNA damage and replication checkpoint kinase in response to exogenous DNA damage. (Right) Rad53 regulates G2/M progression during the cell cycle by restraining mitochondrial function and limiting ROS accumulation.

*S. cerevisiae* has evolved for rapid growth in high glucose, and repress their mitochondria^45^, much like cancer cells. Strikingly, the loss of Rad53 (*rad53Δ* cells) result in a unique, pronounced metabolic rewiring characterized by increased flux through the mitochondrial TCA cycle, and mitochondrial de-repression despite the presence of glucose. This is observed as a selective remodeling of the mitochondrial proteome in *rad53Δ* cells, with non-uniform increases in mitochondrial proteins. This results in a reallocation of carbon flux, towards a higher TCA cycle, and ETC activity in *rad53Δ* cells leading to high membrane potential. However, this altered mitochondrial state in *rad53Δ* cells does not have corresponding increases in OCR and mitochondrial ATP levels, leading to substantial ROS generation and oxidative stress. This role of Rad53 in restraining mitochondrial outputs and resultant ROS therefore places a critical brake on normal G2/M cell cycle progression. Importantly, reducing this high ROS using antioxidants is sufficient to rescue the cell division defects, cell size heterogeneity and delayed G2/M progression in *rad53Δ* cells (Fig 6). All of this reiterates the critical roles of increased ROS and oxidative stress as a constraint for cell-cycle timing^46^ particularly delaying the G2/M transition^47,48^. Restricting ROS is a critical requirement within cells in order to maintain genomic integrity and normal cell proliferation. Our studies thus suggest a likely primordial role for the Rad53 protein in restraining ROS, by placing a check on mitochondrial outputs during fermentative growth. This reveals how a classic ‘check-point’ kinase (Rad53/Chk2) might mediate cell cycle progression by restraining untimely mitochondrial outputs. Mutations in Chk2 are widely observed in a variety of cancers^49–51^, which are themselves associated with high ROS. It will be therefore particularly interesting to explore possible conserved functions of Chk2 in this context. Further, the role of Rad53 in restraining mitochondrial and ROS-outputs are catalytic activity dependent, but distinct from the DNA damage-induced *RAD53*-*SML1* checkpoint axis (Fig 5). Interestingly, recent studies also suggest that Rad53 can function distinctly from the canonical Mec1-Rad53-Sml1 axis^24,26^. An important study implicated a role for Rad53 in maintaining acetyl-CoA pools, which in turn alters gene expression states^24^. This would be consistent with likely consequences of altered mitochondrial function and outputs. These diverse findings can now be addressed through the role of Rad53 uncovered in this study, in restraining mitochondrial outputs and functions, without which cells will substantially alter their carbon allocations. Importantly, these findings with Rad53 relied on the use of fully prototrophic yeast - the natural state of yeasts which are capable of *de novo* amino acid and nucleotide synthesis. Prior studies of Rad53 utilize metabolically deficient, auxotrophic laboratory strains, which are distinct from prototrophic cells^52^. Gene essentiality is conditional, emerging from the interaction between a gene, its genetic background, and the metabolic environment^53,54^, and critical metabolic roles of Rad53 could therefore easily have been missed.

Collectively, our data reveal that beyond critical roles in responses to acute genomic insults, Rad53 has distinct, constitutive roles in restraining mitochondrial activity and maintaining redox homeostasis required for G2/M progression. Importantly, our study reiterates the critical coupling between metabolic state and cell cycle progression, and the need for these processes to be coordinately regulated. Seminal discoveries made using the well-known yeast ultradian metabolic oscillations (YMC), revealed a temporal compartmentalization of metabolic processes during cell division^5,6,14^. These studies established that the DNA replication and mitotic cell division processes are restricted to a reductive phase, and are temporally separated from biomass building processes that occur in oxidative phases with higher mitochondrial activity, thereby allowing cells to protect their genome from oxidative damage (from excessive mitochondrial activity)^12,55^. A loss of Rad53 uncouples the cell division cycle from the YMC^12,56^. Our findings establish that the Rad53 checkpoint kinase itself functions to couple mitochondrial outputs and ROS with appropriate cell cycle progression.

Speculatively, could a primordial function of Rad53 be to restrain ROS during cell cycle progression and growth, and which was later co-opted for the DNA-damage response system? The importance of selectively restraining mitochondrial activity for cell cycle progression is clear, and Rad53 might have evolved to enable cells to maintain a reduced state, permissive for mitotic cell division. Our study therefore opens up exciting new possibilities to address critical homeostatic roles of Rad53 in mitochondrial regulation, and to understand how cells couple redox homeostasis with cell cycle progression.

## Materials and methods

Also see S1 (Appendix I) for extended materials and methods.

### Statistics and graphing

Statistical significance was assessed using two-tailed Welch’s *t*-tests, unless indicated otherwise. All experiments were performed with at least three biological replicates, as specified in the corresponding figure legends. Graphs were visualised using GraphPad Prism (v9.0.1 and v10.5.0) and RStudio (v2025.05.1+513). All the figures, schematics and models were created on Adobe illustrator v30.8.1.

### Yeast strains, media and growth conditions

The prototrophic CEN.PK strain of *Saccharomyces cerevisiae* was used as the wild type in all the experiments. All the strains used in this study are listed in Table S1. Media compositions and growth conditions are described in Extended methods (Appendix 1).

### Microscopic Analysis for budding time and cell size

WT and *rad53Δ* cells were placed on agarose pads and time-lapse imaging was done using a NIKON ECLIPSE Ti2 inverted microscope equipped with a 60× oil-immersion objective. Image analysis was done using FIJI. For automated segmentation, images were processed using Cell-ACDC (v1.7.1) and Statistical analysis and visualization were performed in GraphPad Prism (v10.5.0) and RStudio (v2025.05.1+513) using the ggplot2 and tidyr packages. Detailed protocol is described in Extended methods (Appendix 1).

### Flow Cytometry Analysis

For DNA content analysis, cells were stained with SYTOX Green and analyzed on BD LSR Fortessa X-20 flow cytometer using a 488nM excitation laser as previously described^57^. Detailed protocol is described in Extended methods (Appendix 1).

### Protein extraction and western blotting

Total protein extraction was done using TCA as previously described^58^. Western blots were quantified using ImageJ. The quantification of the blots was done using ImageJ. The statistical significance was determined using Student’s t-test (using GraphPad Prism) Detailed protein extraction and western blotting protocol is described in Extended methods (Appendix 1).

### Metabolite Extraction and analysis

The metabolite extraction and analysis for steady state and metabolic flux measurements were carried out following protocols described previously^29,32^. The metabolites were detected using ABSciex QTRAP 5500 mass spectrometer. The data was acquired using Analyst 1.6.2 software (Sciex) and analyzed using MultiQuant version 3.0.1 (Sciex). Analyzed data was plotted using GraphPad Prism. Raw data and LC-MS/MS parameters are listed in Table S2. Detailed metabolite extraction protocol is described in Extended methods (Appendix 1).

### Nonyl acridine orange (NAO) staining

1 OD_600_. of log phase growing cells were collected, washed once and resuspended in the 1 ml fresh media, YPD (2%). To this 25 μM of Nonyl acridine orange is added and incubated for 15 mins. Following this, the cells are washed once with media, and twice with PBS, resuspended in PBS. The cells were analysed on Attune NxT cytometer using 488 excitation laser. The fluorescence of the cells were analysed FlowJo, and median values were compared.

### Mitotracker and DCFDA fluorescence assays

Mitotracker fluorescence assay was performed as described earlier using MitoTracker Red CMXRos (Invitrogen; M7512). DCFDA assay was performed using CM-H2DCFDA (Invitrogen; C6827). The fluorescence (Ex/Em 579/600 for mitotracker and 495/522 for CM-H2DCFDA) was measured using Thermo Varioscan LUX multimode plate reader. Fluorescence values were normalized to cell number based on absorbance at 600 nm. Detailed protocol is described in extended methods (Appendix 1).

### Oxygen Consumption Rate

Oxygen consumption rate was measured using Agilent Seahorse XFe24 analyzer. The obtained OCR values were normalised to cell number using OD_600_. Details are described in extended methods (Appendix 1).

### ATP measurements

ATP levels were measured using ATP Determination Kit (Invitrogen; A22066) The reaction mixture for ATP measurements were prepared as per manufacturer’s instructions and luminescence was measured using Sirius luminometer (Tiertek Berthold). The relative ATP concentrations were calculated, graphs were plotted and using GraphPad prism 9.0.1. Detailed sample preparation protocol is described in extended methods (Appendix 1).

### SOD activity assay

SOD activity was measured using a Superoxide Dismutase Assay Kit (Cayman Chemicals; 706002) as per manufacturer’s instructions. The relative SOD activity were calculated and plotted using GraphPad prism 9.0.1. Detailed protocol is described in extended methods (Appendix 1).

### Mitochondrial enrichment and proteomics

Mitochondrial enrichment was performed as previously described.^35,36^. Details are described in extended methods (Appendix 1). Proteins were extracted from enriched mitochondrial fractions and analyzed on Thermo EASY-nLC 1200nano System coupled to a Thermo Scientific Orbitrap Fusion Tribrid Mass Spectrometer. The experiment was performed in 3 biological replicates. Data analysis was performed using DIA-NN (v1.8.1). Detailed protocol is described in extended methods (Appendix 1).

---

**Table S1.** List of Strains used in this study.

| **No.** | **Strain name** | **Genotype** | **Source** |
| --- | --- | --- | --- |
| **1** | CEN.PK<br>(*wild-type*) | *Mat a* | Van Dijken et al, 2000 |
| **3** | Rad53-HA | *Mat a RAD53-HA:: kanMX* | This study |
| **4** | Rad53-HA/ *bar1*Δ | *Mat a RAD53-HA::kanMX: bar1Δ:: NAT* | This study |
| **5** | *RAD53/ rad53*Δ | *Mat a/α RAD53/ rad53Δ:: Hyg* | This study |
| **6** | *rad53*Δ | *Mat a rad53Δ:: Hyg* | This study; |
| **7** | Rad53<sup>KD</sup> | *Mat a RAD53<sup>mut</sup>:: Hyg* | This study |
| **7** | *SML1/ sml1*Δ | *Mat a SML1/ sml1Δ:: kanMX* | This study |
| **8** | *sml1*Δ | *Mat a sml1Δ:: kanMX* | This study |
| **9** | *rad53*Δ/ *RAD53:: sml1*Δ/ *SML1* | *Mat a/α Rad53/ rad53Δ:: Hyg: SML1/ sml1Δ:: kanMX* | This study |
| **10** | *rad53*Δ/*sml1*Δ | *Mat a rad53Δ:: Hyg: sml1Δ:: kanMX* | This study |

## Notes

### Competing Interest Statement

The authors have declared no competing interest.

## References

1. Elledge, S. J. Cell cycle checkpoints: preventing an identity crisis. Science 274, 1664–72 (1996).

2. Longhese, M. P., Foiani, M., Muzi-Falconi, M., Lucchini, G. & Plevani, P. DNA damage checkpoint in budding yeast. EMBO Journal 17, 5525–5528 (1998).

3. Ewald, J. C., Kuehne, A., Zamboni, N. & Skotheim, J. M. The Yeast Cyclin-Dependent Kinase Routes Carbon Fluxes to Fuel Cell Cycle Progression. Mol. Cell 62, 532–545 (2016).

4. Patton, E. E. SCFMet30-mediated control of the transcriptional activator Met4 is required for the G1-S transition. EMBO J. 19, 1613–1624 (2000).

5. Tu, B. P., Kudlicki, A., Rowicka, M. & McKnight, S. L. Logic of the Yeast Metabolic Cycle: Temporal Compartmentalization of Cellular Processes. Science (1979). 310, 1152–1158 (2005).

6. Slavov, N. & Botstein, D. Coupling among growth rate response, metabolic cycle, and cell division cycle in yeast. Mol. Biol. Cell 22, 1997–2009 (2011).

7. Ewald, J. C. How yeast coordinates metabolism, growth and division. Curr. Opin. Microbiol. 45, 1–7 (2018).

8. Cai, L., Sutter, B. M., Li, B. & Tu, B. P. Acetyl-CoA induces cell growth and proliferation by promoting the acetylation of histones at growth genes. Mol. Cell 42, 426–37 (2011).

9. Shi, L. & Tu, B. P. Acetyl-CoA induces transcription of the key G1 cyclin CLN3 to promote entry into the cell division cycle in Saccharomyces cerevisiae. Proceedings of the National Academy of Sciences 110, 7318–7323 (2013).

10. Kurat, C. F. et al. Cdk1/Cdc28-Dependent Activation of the Major Triacylglycerol Lipase Tgl4 in Yeast Links Lipolysis to Cell-Cycle Progression. Mol. Cell 33, 53– 63 (2009).

11. Muller, E. G. Thioredoxin deficiency in yeast prolongs S phase and shortens the G1 interval of the cell cycle. J. Biol. Chem. 266, 9194–202 (1991).

12. Chen, Z., Odstrcil, E. A., Tu, B. P. & McKnight, S. L. Restriction of DNA replication to the reductive phase of the metabolic cycle protects genome integrity. Science 316, 1916–9 (2007).

13. López-Grueso, M. J. et al. Knockout of PRDX6 induces mitochondrial dysfunction and cell cycle arrest at G2/M in HepG2 hepatocarcinoma cells. Redox Biol. 37, 101737 (2020).

14. Tu, B. P. et al. Cyclic changes in metabolic state during the life of a yeast cell. Proc. Natl. Acad. Sci. U. S. A. 10.1073/pnas.0708365104 (2007) doi:10.1073/pnas.0708365104.

15. Chiu, J. et al. Cell cycle sensing of oxidative stress in Saccharomyces cerevisiae by oxidation of a specific cysteine residue in the transcription factor Swi6p. J. Biol. Chem. 286, 5204–14 (2011).

16. Weinert, T. A., Kiser, G. L. & Hartwell, L. H. Mitotic checkpoint genes in budding yeast and the dependence of mitosis on DNA replication and repair. Genes Dev. 10.1101/gad.8.6.652 (1994) doi:10.1101/gad.8.6.652.

17. Allen, J. B., Zhou, Z., Siede, W., Friedberg, E. C. & Elledge, S. J. The SAD1/RAD53 protein kinase controls multiple checkpoints and DNA damage-induced transcription in yeast. Genes Dev. 8, 2401–2415 (1994).

18. Pellicioli, A. & Foiani, M. Signal Transduction: How Rad53 Kinase Is Activated. Current Biology 15, R769–R771 (2005).

19. Branzei, D. & Foiani, M. The Rad53 signal transduction pathway: Replication fork stabilization, DNA repair, and adaptation. Exp. Cell Res. 312, 2654–9 (2006).

20. Hoch, N. C. et al. Molecular basis of the essential s phase function of the rad53 checkpoint kinase. Mol. Cell. Biol. 33, 3202–13 (2013).

21. Zhao, X., Muller, E. G. D. & Rothstein, R. A suppressor of two essential checkpoint genes identifies a novel protein that negatively affects dNTP pools. Mol. Cell 2, 329–340 (1998).

22. Koc, A. & Merrill, G. F. Checkpoint deficient rad53-11 yeast cannot accumulate dNTPs in response to DNA damage. Biochem. Biophys. Res. Commun. 10.1016/j.bbrc.2006.12.049(2007) doi:10.1016/j.bbrc.2006.12.049.

23. Covo, S., Westmoreland, J. W., Reddy, A. K., Gordenin, D. A. & Resnick, M. A. RAD53 is limiting in double-strand break repair and in protection against toxicity associated with ribonucleotide reductase inhibition. DNA Repair (Amst). 11, 317– 23 (2012).

24. Bruhn, C. et al. The Rad53CHK1/CHK2-Spt21NPAT and Tel1ATM axes couple glucose tolerance to histone dosage and subtelomeric silencing. Nat. Commun. 11, 4154 (2020).

25. Bu, P. et al. DNA damage response activates respiration and thereby enlarges dNTP pools to promote cell survival in budding yeast. Journal of Biological Chemistry 294, 9771–9786 (2019).

26. Ho, B. et al. Mec1-independent activation of the Rad53 checkpoint kinase revealed by quantitative analysis of protein localization dynamics. Elife 12, (2023).

27. Zhao, X. et al. Mutational and structural analyses of the ribonucleotide reductase inhibitor Sml1 define its Rnr1 interaction domain whose inactivation allows suppression of mec1 and rad53 lethality. Mol. Cell. Biol. 20, 9076–83 (2000).

28. Padovani, F., Mairhörmann, B., Falter-Braun, P., Lengefeld, J. & Schmoller, K. M. Segmentation, tracking and cell cycle analysis of live-cell imaging data with Cell-ACDC. BMC Biol. 20, 174 (2022).

29. Walvekar, A., Rashida, Z., Maddali, H. & Laxman, S. A versatile LC-MS/MS approach for comprehensive, quantitative analysis of central metabolic pathways. Wellcome Open Res. 3, 122 (2018).

30. Walsh, C. T., Tu, B. P. & Tang, Y. Eight Kinetically Stable but Thermodynamically Activated Molecules that Power Cell Metabolism. Chem. Rev. 118, 1460–1494 (2018).

31. Vengayil, V., Niphadkar, S., Adhikary, S., Varahan, S. & Laxman, S. The deubiquitinase Ubp3/Usp10 constrains glucose-mediated mitochondrial repression via phosphate budgeting. Elife 12, (2024).

32. Niphadkar, S., Sreedharan, S., Vengayil, V. & Laxman, S. Protocol to quantitatively assess glycolysis and related carbon metabolic fluxes using stable isotope tracing in Crabtree-positive yeasts. STAR Protoc. 6, 103786 (2025).

33. Rashida, Z. & Laxman, S. The pentose phosphate pathway and organization of metabolic networks enabling growth programs. Curr. Opin. Syst. Biol. 28, 100390 (2021).

34. Mileykovskaya, E. et al. Cardiolipin binds nonyl acridine orange by aggregating the dye at exposed hydrophobic domains on bilayer surfaces. FEBS Lett. 507, 187–90 (2001).

35. Adhikary, S., Vengayil, V. & Laxman, S. Isolation of Mitochondria From Yeast to Estimate Mitochondrial Pools of Inorganic Phosphate. Bio. Protoc. 15, e5370 (2025).

36. Liao, P.-C., Boldogh, I. R., Siegmund, S. E., Freyberg, Z. & Pon, L. A. Isolation of mitochondria from Saccharomyces cerevisiae using magnetic bead affinity purification. PLoS One 13, e0196632 (2018).

37. Demichev, V., Messner, C. B., Vernardis, S. I., Lilley, K. S. & Ralser, M. DIA-NN: neural networks and interference correction enable deep proteome coverage in high throughput. Nat. Methods 17, 41–44 (2020).

38. Vögtle, F.-N. et al. Landscape of submitochondrial protein distribution. Nat. Commun. 8, 290 (2017).

39. Morgenstern, M. et al. Definition of a High-Confidence Mitochondrial Proteome at Quantitative Scale. Cell Rep. 19, 2836–2852 (2017).

40. Di Bartolomeo, F. et al. Absolute yeast mitochondrial proteome quantification reveals trade-off between biosynthesis and energy generation during diauxic shift. Proc. Natl. Acad. Sci. U. S. A. 117, 7524–7535 (2020).

41. Singh, G. & Laxman, S. Lighting up mitochondrial membranes: Reading and writing the physical state of the inner mitochondrial membrane. Curr. Opin. Chem. Biol. 94, 102734 (2026).

42. Singh, G., Vengayil, V., Khanna, A., Adhikary, S. & Laxman, S. Active control of mitochondrial network morphology by metabolism-driven redox state. Proc. Natl. Acad. Sci. U. S. A. 122, e2421953122 (2025).

43. Jastroch, M., Divakaruni, A. S., Mookerjee, S., Treberg, J. R. & Brand, M. D. Mitochondrial proton and electron leaks. Essays Biochem. 47, 53–67 (2010).

44. Saffi, J., Sonego, L., Varela, Q. D. & Salvador, M. Antioxidant activity of L-ascorbic acid in wild-type and superoxide dismutase deficient strains of Saccharomyces cerevisiae. Redox Rep. 11, 179–84 (2006).

45. Diaz-Ruiz, R., Rigoulet, M. & Devin, A. The Warburg and Crabtree effects: On the origin of cancer cell energy metabolism and of yeast glucose repression. Biochimica et Biophysica Acta - Bioenergetics Preprint at 10.1016/j.bbabio.2010.08.010 (2011).

46. de Souza Pereira, R. & Geibel, J. Direct observation of oxidative stress on the cell wall of Saccharomyces cerevisiae strains with atomic force microscopy. Mol. Cell. Biochem. 201, 17–24 (1999).

47. Zhang, Y. et al. Oxidative stress-induced DNA damage of mouse zygotes triggers G2/M checkpoint and phosphorylates Cdc25 and Cdc2. Cell Stress Chaperones 21, 687–96 (2016).

48. He, P. et al. AMPK Activity Contributes to G2 Arrest and DNA Damage Decrease via p53/p21 Pathways in Oxidatively Damaged Mouse Zygotes. Front. Cell Dev. Biol. 8, 539485 (2020).

49. Bartek, J., Falck, J. & Lukas, J. CHK2 kinase--a busy messenger. Nat. Rev. Mol. Cell Biol. 2, 877–86 (2001).

50. Bai, M. et al. ROS-ATM-CHK2 axis stabilizes HIF-1α and promotes tumor angiogenesis in hypoxic microenvironment. Oncogene 44, 1609–1619 (2025).

51. Lulli, M. et al. DNA Damage Response Protein CHK2 Regulates Metabolism in Liver Cancer. Cancer Res. 81, 2861–2873 (2021).

52. Pronk, J. T. Auxotrophic yeast strains in fundamental and applied research. Appl. Environ. Microbiol. 68, 2095–100 (2002).

53. Bosch-Guiteras, N. & van Leeuwen, J. Exploring conditional gene essentiality through systems genetics approaches in yeast. Curr. Opin. Genet. Dev. 76, 101963 (2022).

54. Kaplan, K., Levkovich, S. A., DeRowe, Y., Gazit, E. & Laor Bar-Yosef, D. Mind your marker: the effect of common auxotrophic markers on complex traits in yeast. FEBS J. 291, 2209–2220 (2024).

55. Yu, F.-X., Dai, R.-P., Goh, S.-R., Zheng, L. & Luo, Y. Logic of a mammalian metabolic cycle: an oscillated NAD+/NADH redox signaling regulates coordinated histone expression and S-phase progression. Cell Cycle 8, 773–9 (2009).

56. Chen, Z. & McKnight, S. L. A conserved DNA damage response pathway responsible for coupling the cell division cycle to the circadian and metabolic cycles. Cell Cycle 6, 2906–12 (2007).

57. Gupta, R. et al. A tRNA modification balances carbon and nitrogen metabolism by regulating phosphate homeostasis. Elife 8, (2019).

58. Gupta, R., Adhikary, S., Dalpatraj, N. & Laxman, S. An economic demand-based framework for prioritization strategies in response to transient amino acid limitations. Nat. Commun. 15, 7254 (2024).

