## Supplementary material for "Rad53/Chk2 couples mitochondrial outputs and ROS with G2/M cell cycle progression": SI

**This PDF file includes:**

Appendix I

Figures S1 to S6

Legends for Movies S1 to S6

Legends for Datasets S2 to S6

SI References

**Other supporting materials for this manuscript include the following:**

Movies S1 to S6

Datasets S2 to S6

### Appendix I

#### Extended methods

##### 1. Media and growth conditions

For cell growth, an overnight preculture was grown at 30°C (OD<sub>600</sub> ~5-6) in YPD medium (1% yeast extract, 2% peptone, 2% dextrose), and subsequently sub-cultured in the same medium (starting OD<sub>600</sub> ~0.2) and grown to an OD<sub>600</sub> ~0.7-0.8. For α-factor G1 arrest – Overnight grown cells were diluted to 0.1 OD<sub>600</sub> and sub-cultured in fresh YPD media. These were grown briefly till OD<sub>600</sub> ~0.2-0.3. Cells with *bar1Δ* were then incubated with 50 ng/mL α-factor for 3 hours at 30°C in shaking incubator as previously described<sup>1</sup>. The cells were then washed with equal volume of YPD once, then with autoclaved water twice, and resuspended in fresh media to release the arrested cells.

##### 2. Sporulation and tetrad Dissection

Diploid parent strains were cultured in YPD overnight. Following this, 200 μL of the primary culture was centrifuged at 1500g for 5 min at RT, and washed with 5 ml sterile water. The pellet was resuspended in 2.5ml of Sporulation medium (3g potassium acetate and 0.2 raffinose in 1L of water) and incubated at 30°C for 2-5 days, following which tetrads were confirmed using a microscope. 500μL of this sporulated culture was washed with 1 ml sterile water. The pellet was resuspended in 50 μL of Zymolyase (0.5mg/ml in 1M Sorbitol) and incubated at 30°C for ~10-12 min, and tetrads were dissected using a dissection microscope (Singer Instruments). The spores were then scored for their mating type-based on PCR analysis<sup>2</sup>.

##### 3. Serial dilution growth assay

Cells were grown overnight in YPD (2%) and sub-cultured to an OD<sub>600</sub> ~ 0.8. Then, 1 OD<sub>600</sub> cells were collected, washed once in water and resuspended in 1 ml of autoclaved water. This cell suspension is then serially diluted by 10-fold. 5 μL of the different diluted suspensions was then spotted on media plates. For H<sub>2</sub>O<sub>2</sub> sensitivity assay, the serial dilution spots are done on media with 1 mM H<sub>2</sub>O<sub>2</sub><sup>3</sup>.

##### 4. Microscopic Analysis for budding time and cell size

Overnight cultures of WT and *rad53Δ* were diluted to 0.2-0.3 O.D<sub>600</sub>. The cells were then collected and resuspended in ~100-150 μL media. 2 μL of these cells were placed on 1% agarose pads with suitable media. The agarose pad was inverted and placed on a glass-bottom confocal dish incubated at 30°C. Time-lapse phase-contrast imaging of the cells was done every 10 minutes over a 6-hour period using a NIKON ECLIPSE Ti2 inverted microscope equipped with a 60× oil-immersion objective. Images were stacked and processed using FIJI. Budding time was scored manually for more than 100 cells per strain under each condition. For automated segmentation, images were processed using Cell-ACDC (v1.7.1) with the YeaZ\_v2 deep learning model, which is optimized for budding yeast segmentation in phase-contrast microscopy<sup>4</sup>. Morphometric parameters — including cell area, volume, circularity, and aspect ratio — were extracted for each segmented cell. To exclude imaging artifacts and segmentation debris, objects with a projected area below 12 μm<sup>2</sup> were removed from analysis, as this falls below the expected average size of a yeast cell (approximate area ~12.5 μm<sup>2</sup> based on a mean yeast diameter of ~4 μm). A minimum of 600 cells per strain were analyzed across all conditions. Segmentation masks and single-cell measurements were exported as CSV files for downstream analysis. Statistical analysis and visualization were performed in GraphPad Prism (v10.5.0) and RStudio (v2025.05.1+513) using the ggplot2 and tidyr packages.

##### 5. Flow Cytometry Analysis

1 O.D<sub>600</sub> (1 × 10<sup>7</sup> cells) of log phase cells were collected and fixed overnight with 70% ethanol at 4°C. The cells after fixing were collected by centrifugation and transferred to a microcentrifuge tube. The cells were treated with RNase for 2-12hrs at 37°C, followed by protease treatment at 37°C as mentioned in (Haase & Reed, 2002). For DNA content analysis, cells were stained with SYTOX Green by resuspending ~50 μL of cell suspension in 1 mL of staining solution. The samples were then sonicated and analyzed on BD LSR Fortessa X-20 flow cytometer using a 488nm excitation laser<sup>1</sup>.

##### 6. Protein extraction and western blotting

For Rad53 protein level estimation, Rad53 was tagged at its chromosomal locus with a C-terminal HA epitope, cultured and log phase cells were collected. For Cox2 and

γH2AX, the cells were pelleted using 10% trichloroacetic acid (TCA). Protein extraction was performed using the TCA extraction method<sup>5</sup>. Briefly, the pelleted cells were resuspended in 400 μl 10 % TCA solution and the cells were lysed by bead beating (3 times, 20 seconds each with 1 min cooling in ice-water slurry between each cycle). Lysate extracts were centrifuged and the protein pellets thus obtained were resuspended in 400 μl SDS-glycerol buffer (7.3% SDS, 29.1% glycerol and 83.3 mM Tris base), and heated for 10 minutes at 100°C. The supernatant was collected after centrifugation (20000g/10 min), and total protein estimation was done using BCA assay (BCA assay kit, G-Biosciences). Samples were normalized (to maintain constant protein amounts) in SDS-glycerol buffer. The samples were run on 4-12 % Bis-Tris gels (Invitrogen). Western blots were developed using the following antibodies – anti-HA raised in mouse (1:2000; Sigma-Aldrich, 11583816001), anti-Cox2 raised in mouse (1:2000; MTCO2 Monoclonal Antibody, Invitrogen, 459150), anti-Histone H2A (phosphor S129) raised in rabbit (1:500; Abcam, ab15083), anti-Porin raised in mouse (1:2000; Invitrogen, 459500), anti-Ildh1 raised in goat (1:2000; Sigma-Aldrich, SAB2501682), anti-mouse horseradish peroxidase (HRP)-conjugated antibody (1:4000; Cell Signaling Technology, 7076S), anti-rabbit HRP-conjugated antibody (1:4000; Cell Signaling Technology, 7074S) and anti-goat HRP-conjugated antibody (1:4000; Jackson ImmunoResearch, C839K62). For developing the blot, a standard enhanced chemiluminescence reagent (WesternBright ECL HRP substrate, Advansta, K12045) was used. Coomassie-stained gels were used as the loading controls. The quantification of the blots was done using ImageJ. The statistical significance was determined using two-tailed unpaired, Welch's T-test (using GraphPad Prism)

### **7. Metabolite Extraction**

The metabolite extraction and analysis were carried out following protocols described previously<sup>6</sup>. For each experiment, 10 OD<sub>600</sub> cells were used for metabolite extraction. First, the cells were quenched for 5 minutes in 60% methanol (maintained at -45°C). After centrifugation, the cell pellet was resuspended in the extraction buffer (75% ethanol) and incubated at 80°C for 5 mins followed by incubation on ice and centrifugation. The supernatant was collected, dried using a vacuum concentrator, and then stored at -80°C till further use. TCA metabolites (with functional carbonyl and carboxyl groups) were derivatized with O-benzylhydroxylamine (OBHA). The metabolite extract was dissolved in 1-ethyl-3-(3-dimethylaminopropyl) carbodiimide (EDC); OBHA was added to this mixture and mixed by shaking for 1 hour, to this, ethyl acetate was added and mixed for additional 10 minutes. The top layer was extracted in a new vial, this is done three times and top layers were pooled together. The derivatized extract was dried using a vacuum concentrator and stored at -80°C till further use. For glutathione measurements, samples were resuspended in 0.1% formic acid and analyzed on the same day of extraction. For TCA intermediates, the extracts were resuspended in a solution of 1:1 methanol: water. Metabolites were separated on Synergi 4-μm Fusion-RP 80 Å (100/150 × 4.6 mm) LC column (Phenomenex, 00F-4424-E0) using the Shimadzu Nexera UHPLC system<sup>6</sup>. Glutathione and TCA intermediates was separated using 0.1% formic acid (solvent A) and methanol and 0.1% Formic acid (solvent B) as solvents. The solvents used for separating sugar phosphatases and trehalose were 5 mM ammonium acetate in water (solvent A) and acetonitrile (solvent B). The metabolites were detected using ABSciex QTRAP 5500 mass spectrometer. The data was acquired using Analyst 1.6.2 software (Sciex) and analyzed using MultiQuant version 3.0.1 (Sciex). Analyzed data was plotted using GraphPad Prism. Raw data and LC-MS/MS parameters are listed in Table S2.

### 8. <sup>13</sup>C<sub>6</sub> glucose based metabolic flux measurements

The carbon flux into TCA cycle was estimated by measuring the relative incorporation of <sup>13</sup>C label into TCA cycle intermediates. Cells were grown in YPD medium (2% glucose) to OD<sub>600</sub> ~0.6 and shifted to fresh medium with 1% glucose. The OD<sub>600</sub> was measured 30 minutes after the shift and cells corresponding to ~5 OD<sub>600</sub> were collected, and <sup>13</sup>C<sub>6</sub> glucose (Cambridge Isotope Laboratories, CLM-1396) was pulsed at a final concentration of 1%, making the total glucose concentration in the medium 2% (1% labelled + 1% unlabelled). Metabolites were extracted as described earlier, 15 minutes after labelled glucose addition<sup>7</sup>. The relative label incorporation was calculated by measuring the area under the curve for m+2 labelled TCA metabolites using MultiQuant version 3.0.1 (Sciex). Analyzed data was plotted using GraphPad Prism.

### 9. Mitotracker fluorescence assays

Cells were grown in YPD (2%) media to mid-log phase (OD<sub>600</sub>~0.6–0.8), harvested by centrifugation, washed, and resuspended in fresh media. MitoTracker Red CMXRos

(Invitrogen; M7512) was added to cells at a final concentration of 200 nM, and incubated for 30 mins at 30 °C. Following staining, cells were washed and resuspended in fresh media, then fixed with formaldehyde for 15-20 mins. Cells were washed twice with PBS and resuspended in 1 ml PBS for analysis. For plate reader measurements, 250 µL of the cell suspension was transferred to 96-well plates, and fluorescence was measured at an excitation wavelength of 579 nm and an emission wavelength of 600 nm. Fluorescence values were normalized to cell number based on absorbance (OD<sub>600</sub>)<sup>3</sup>.

### **10. Oxygen Consumption Rate**

1 ml of the XF Calibrant solution was aliquoted into each well of the utility plate and the sensor cartridge plate was hydrated overnight as per the manufacturer's instructions. The sensor cartridge was loaded with complex IV inhibitor sodium azide and equilibrated in the Seahorse XFe24 analyzer one hour prior to the start of the assay. The culture plate was coated with 50 µl poly-L-lysine and incubated for 1 hour at RT. Excess poly-L-lysine was removed and the plate dried at 30°C for 30 minutes. The cells were grown in standard high glucose to an OD<sub>600</sub>~0.6 and collected, washed and resuspended in fresh media. The samples were aliquoted to poly-L-lysine coated wells at a final cell number in each well of ~3x10<sup>5</sup> cells. The plate was then centrifuged for 2 min at 100xg (acceleration 2, brake 2) and incubated at 30°C for 30 minutes. The plate was loaded to the Seahorse XFe24 analyzer and basal OCR measured over time. A minimum of three measurements were taken with intermittent 2 min mixing and waiting steps. This was followed by sodium azide injection to the sample wells, and three measurements were taken with intermittent 2 min mixing and waiting steps. The obtained values were then normalised to cell OD<sub>600</sub> measured at the end of the experiment.

### **11. Sample preparation for ATP measurements**

Cells were grown in high glucose and at OD<sub>600</sub> ~0.8, 10 OD<sub>600</sub> were pelleted at 4°C. The pellet was treated with 300 µl ice-cold TCA (5%), resuspended and incubated in ice for 15 mins. The suspension was diluted such that the final TCA concentration is 0.1% using 20 mM Tris-HCl, Ph 7.0. For calculating the contribution of mitochondria to total ATP, cells were grown in high glucose and at OD<sub>600</sub> ~0.6, sodium azide was added at a final concentration of 1 mM and incubated for 45 minutes at 30°C in a shaking incubator. The ATP estimation was done by ATP Determination Kit (Invitrogen; A22066). The reaction mixture for ATP measurements were prepared as per manufacturer's instructions and luminescence was measured using Sirius luminometer (Tiertek Berthold). The relative ATP concentrations were calculated, graphs were plotted and using GraphPad prism 9.0.1.

### **12. DCFDA fluorescence assay**

Cells were grown in YPD (2%) media to mid-log phase ($OD_{600} \sim 0.6-0.8$), harvested by centrifugation, washed, and resuspended in fresh media. CM-H2DCFDA (Invitrogen; C6827) was added to cells at a final concentration of $10 \mu\text{M}$, and incubated for 30 mins at $30^\circ\text{C}$. Cells were then washed and resuspended in fresh media, then fixed with formaldehyde for 15-20 mins. Cells were washed twice with PBS and resuspended in 1 ml PBS for analysis. For plate reader measurements, $200 \mu\text{L}$ of the cell suspension was transferred to 96-well plates, and fluorescence was measured at an excitation wavelength of 495 nm and an emission wavelength of $522 \text{ nm}^8$. Fluorescence values were normalized to cell number based on absorbance ($OD_{600}$).

### **13. Sample preparation for SOD activity assay**

Cells were grown in YPD (2%) media to mid-log phase ($OD_{600} \sim 0.6-0.8$), harvested by centrifugation at $1500g$ for 10 minutes at $4^\circ\text{C}$. Cells Lysis (20 mM HEPES, pH 7.2, 1 mM EDTA, 50% Glycerol, KCl) was done by 1 min bead beating, 1 min on ice, for 2 times. Protease inhibitors are not used in the buffer as it interferes with the assay. The lysed cells were centrifuged at $15000 g$ at $4^\circ\text{C}$ for 5 minutes, supernatant (protein) was collected and stored on ice until use. The total protein in lysed cells was determined by BCA assay with a BSA standard. $1 \mu\text{g}$ of protein/sample was used in the assay. SOD activity was measured using a Superoxide Dismutase Assay Kit (Cayman Chemicals; 706002). The SOD standards and the assay mixture was prepared as per manufacturer's instruction; transferred to a 96-well plates and absorbance was measured at 455 nm. The calculations were done as per the instructions in the Kit, and relative SOD activity were calculated and plotted using GraphPad prism 9.0.1.

### **14. Ascorbic acid recovery**

For DCFDA fluorescence, cells were resuspended in fresh YPD and YPD with Ascorbic Acid (100 mM), then stained with CM-H<sub>2</sub>DCFDA. The samples were then processed and analysed as mentioned in (13). For microscopic analysis, the cells were placed on 1% agarose pad of YPD with Ascorbic Acid (100 mM) and imaged and analysed as mentioned in (4). For growth assays, the overnight grown cells are sub-cultured in YPD and YPD with Ascorbic Acid (100 mM) and growth is measured over 18 hours.

### **15. Mitochondrial enrichment**

Mitochondrial enrichment was performed as described in literature<sup>9</sup>, with additional standardization as needed<sup>3,10</sup>. ~200 OD of WT and rad53 cells were collected by centrifugation at 2000g for 5 min at 4°C. The pellet is washed in cold water by centrifugation at 1500g, 5 min at 4°C and incubated in Tris-DTT buffer (0.1 M Tris base, 10 mM DTT, pH 9.4) (5 ml/g pellet weight). This is followed by incubation with zymolyase 20T (0832092, MP biomedical) (5 mL/g) in SEH for 60 minutes, 30°C in a shaking incubator. Resulting spheroplasts are then collected by centrifugation and washed. The pellet is then resuspended in ice-cold SEH buffer with protease inhibitors (1 mM PMSF, 0.5 mg/ml pepstatin A and 0.5 µg/ml leupeptin) (5 mL/g). The resuspended spheroplasts are homogenised using a Dounce homogeniser. The homogenised suspension was centrifuged at 1500 g for 5 mins and supernatant were collected. The supernatant was then centrifuged at a higher speed of 14000 g for 10 mins at 4°C. The pellet is resuspended in SEH buffer (5 mL/g pellet weight) and centrifuged at 700g x 5', collected supernatant and again centrifuged at a higher speed, 14000 g, 10 mins at 4°C. The pellets were resuspended in 700 µL ice-cold SEH buffer with protease inhibitor to obtain the mitochondria-enriched fraction (MEF).

### **16. Sample preparation for Proteomics**

The mitochondrial (MEF) pellet is resuspended in 10% TCA and lysed by bead beating for 30 sec, twice, 1 min on ice in between. Protein extraction was done as mentioned above. The total protein amounts were estimated using BCA assay (BCA assay kit, G-Biosciences). Around 100 µg of protein per sample was then ran on a 4-12% Bis-Tris gels (Invitrogen). The protein bands are cut from the gel and washed with TEAB, chopped and incubated with dithiothreitol, followed by incubation with iodoacetamide. The alkylated proteins were digested by Trypsin (G biosciences) for 12 hrs at 37°C. Trypsin-digested peptides were dissolved in 0.1% formic acid and were analyzed on Thermo EASY-nLC 1200nano System coupled to a Thermo Scientific Orbitrap Fusion Tribrid Mass Spectrometer. The experiment was performed in 3 biological replicates.

### **17. LC-MS/MS acquisition**

Peptides were separated using an EASY-nLC system coupled to an Orbitrap Fusion mass spectrometer operated in positive ion mode and data-independent acquisition (DIA) mode. Peptides were loaded onto the analytical column and separated at a flow rate of 300 nL/min using a 95-min gradient from 10% to 35% solvent B (acetonitrile containing 0.1% formic acid), followed by column washing and re-equilibration, resulting in a total acquisition time of 125 min per sample. Full MS scans were acquired in the Orbitrap at a resolution of 120,000 over an m/z range of 350–2000. DIA MS/MS scans were acquired at a resolution of 30,000 using 25 m/z isolation windows across a precursor mass range of 450–1200 m/z with HCD fragmentation at 30% collision energy.

### **18. Proteomics data analysis**

Data-independent acquisition (DIA) raw files acquired on an Orbitrap mass spectrometer were analysed using DIA-NN (v1.8.1) in library-free (directDIA) mode. A reviewed *Saccharomyces cerevisiae* reference proteome (UniProt ID: UP000002311, downloaded February 2024) was used for analysis. Protein digestion was specified as trypsin (cleavage at K/R) allowing up to one missed cleavage, with a peptide length cut-off of 7–30 amino acids. Carbamidomethylation of cysteine was set as a fixed modification, and methionine oxidation as a variable modification (maximum two variable modifications per peptide). Precursor and fragment ion m/z ranges were set to 300–1800 and 200–1800, respectively, with precursor charge states from +1 to +4. False discovery rate (FDR) was controlled at 1% at both precursor and protein levels. Quantification was performed using fragment ion intensities, and protein group intensities were used for downstream analysis within DIA-NN.

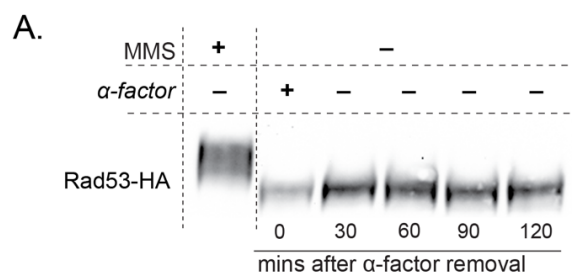

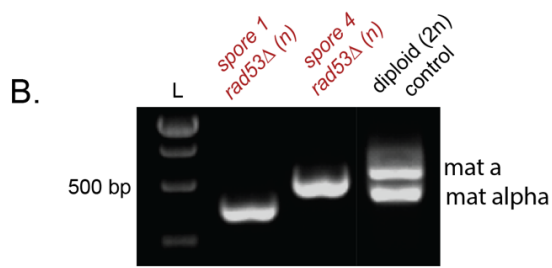

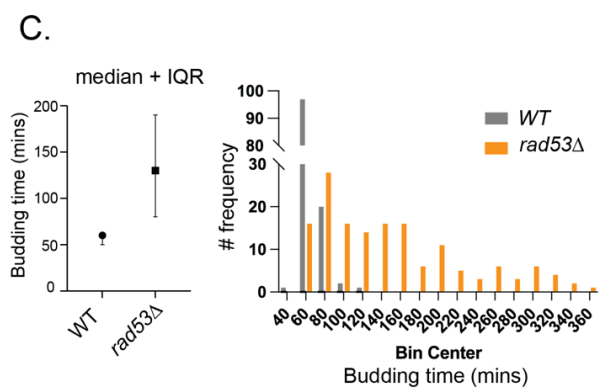

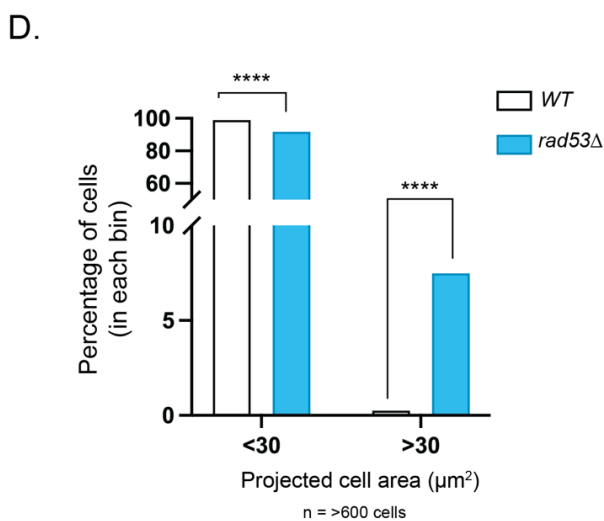

**E.**

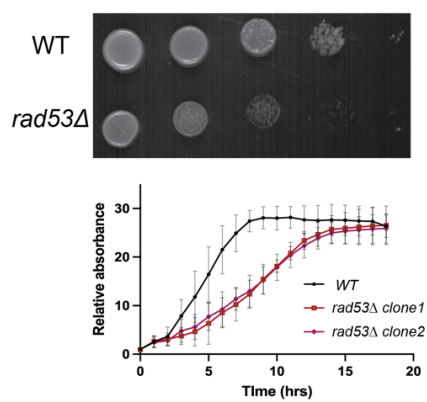

**F.**

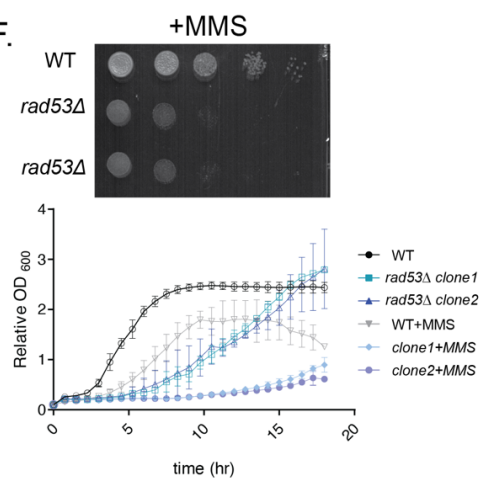

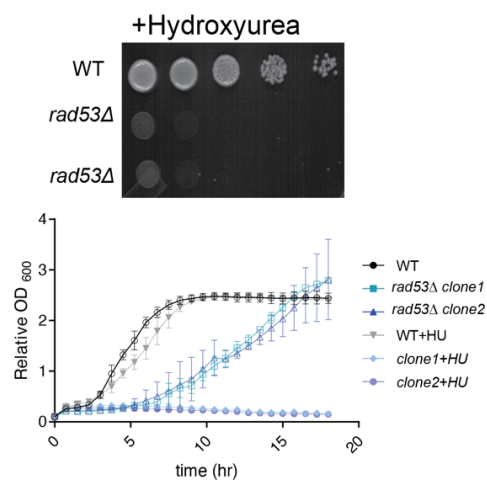

285

286

287

288

**Fig S1.**

**A.** Rad53 protein levels and electrophoretic mobility on an SDS-PAGE gel across cell cycle. Rad53 level was monitored over time by western blotting following release from α-factor-induced G1 arrest into fresh medium.

**B.** PCR based determination of mating type of newly generated *rad53Δ* daughter spores in Fig 1C.

**C.** (Left) Median budding time and interquartile range (IQR) of WT and *rad53Δ* cells. WT cells exhibited a median budding time of 60 min (IQR, 50–60 min), whereas *rad53Δ* cells showed a prolonged budding time with a median of 130 min (IQR, 80–190 min). (Right) Frequency distribution of budding times (right), generated by grouping budding times into 20-min intervals. WT cells (grey) exhibit a narrow distribution centered around 60 min, whereas *rad53Δ* cells (orange) display a broader distribution with a pronounced shift toward longer budding times, indicative of increased cell-to-cell variability. (WT $n = 120$; *rad53Δ* $n = 150$, three independent experiments)

**D.** Projected cell areas of WT and *rad53Δ* cells quantified from segmented images and binned into cells with projected areas of $<30 \mu\text{m}^2$ or $>30 \mu\text{m}^2$. *rad53Δ* cells contained a significantly enriched subpopulation of enlarged cells ($>30 \mu\text{m}^2$) compared with WT. Statistical significance was determined using Fisher's exact test (\*\*\*\*$p < 0.0001$, odds ratio = 27.6; WT, $n = 600$; *rad53Δ*, $n = 700$, pooled from three independent experiments).

**E.** (Up) Serial dilution-based growth assays on solid agar media (YPD), which shows slower growth in *rad53Δ* cells. (Down) Independent clones of *rad53Δ* cells consistently show slower growth in liquid growth experiment DNA damage response. Data are displayed as means ± SD, $n = 3$ (per time point).

**F.** Growth of WT and *rad53Δ* cells following exogenous DNA damage induced by methyl methanesulfonate (MMS) and hydroxyurea (HU). Sensitivity to MMS and HU were assessed by serial dilution spot assays and growth curves, respectively.

A. Steady state central carbon metabolites

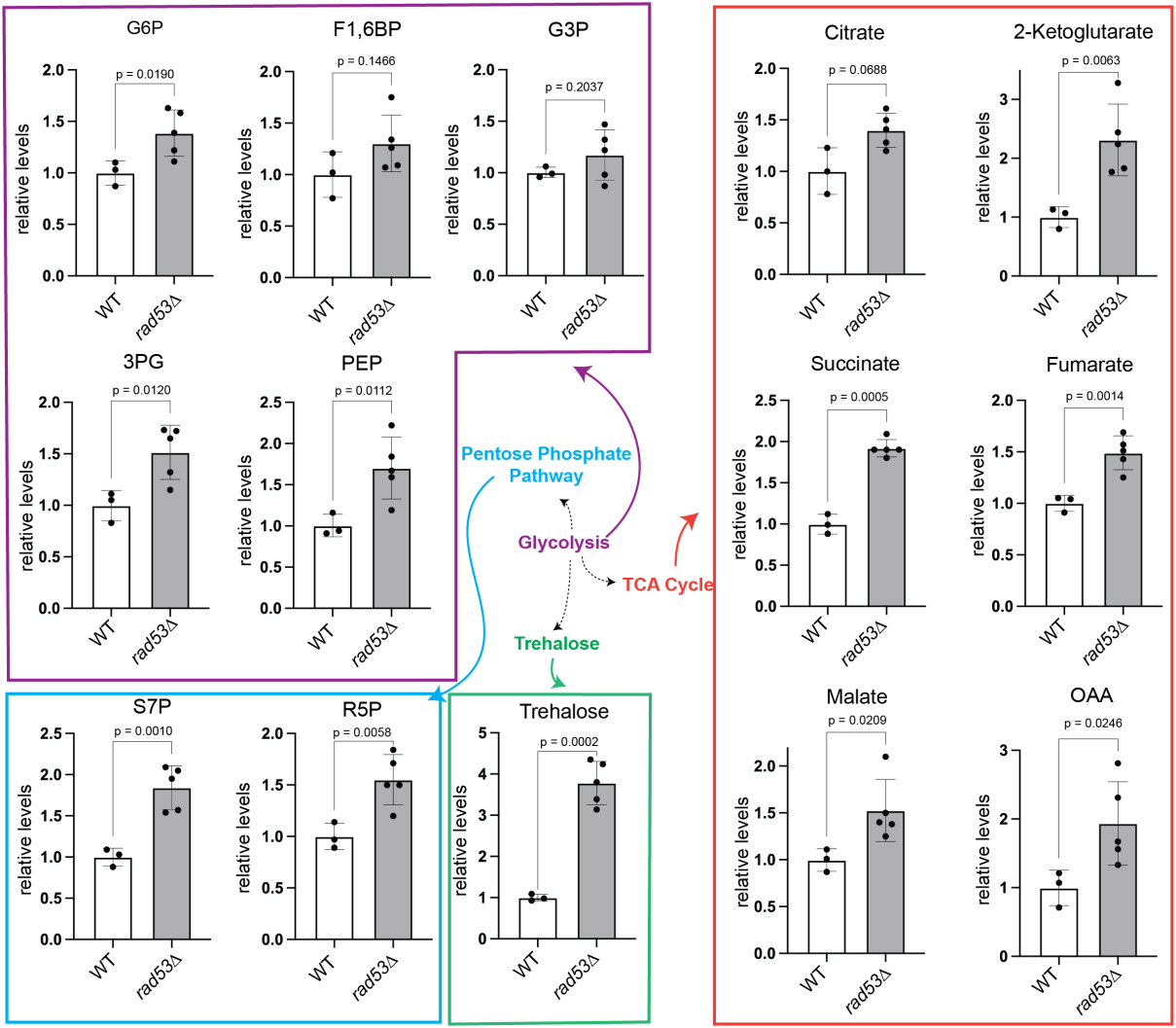

B. Carbon utilization from glucose/ carbon flux

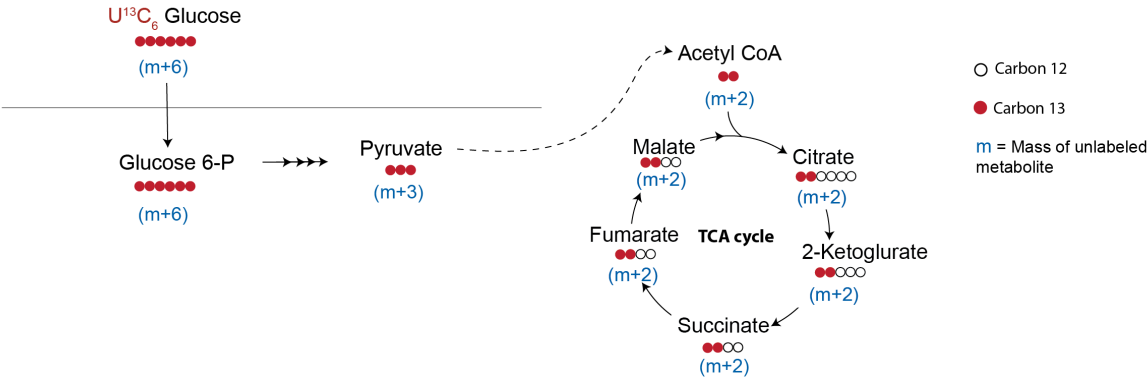

322 **Fig S2.**

323 **A.** Relative steady state levels of central carbon metabolites of *rad53Δ* and WT cells
324 as measured using quantitative LC/MS/MS, shown as bar plots, corresponding to the
325 heat map presented in Fig. 2A. Data are displayed as means ± SD, $n \geq 3$. \*$p < 0.05$,
326 \*\*$p < 0.01$, \*\*\*$p < 0.001$, \*\*\*\*$p < 0.0001$, two-tailed, unpaired Welch's T test.

327 **B.** Schematic illustrating the carbon allocation from labeled $^{13}\text{C}_6$ glucose into
328 tricarboxylic acid (TCA) cycle intermediates (and appropriate mass increase).

329

330

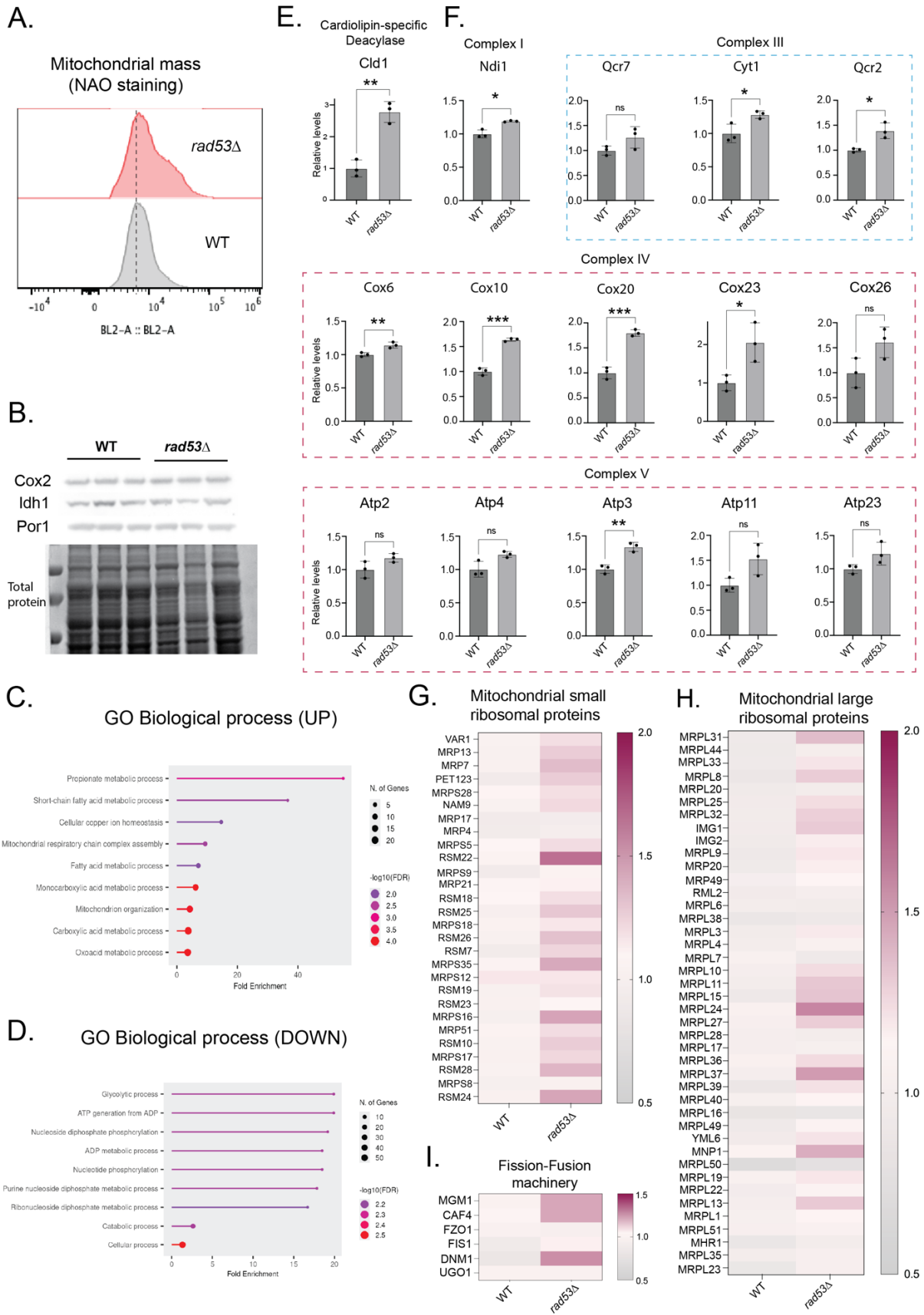

**Fig S3.**

**A.** Distribution of NAO-stained WT and *rad53Δ* cells based on flow cytometry analysis. *rad53Δ* cells exhibited higher median NAO fluorescence compared with WT, indicating elevated cardiolipin-associated fluorescence. Data displayed as means ± SD , n = 4. \*\*p<0.01, two-tailed unpaired Welch's T test.

**B.** Input controls depicting amounts of total isolated mitochondrial proteins that were injected for the biological replicates of *rad53Δ* and WT cells. Mitochondrial proteins Cox2, Por1, and Idh1 was evaluated by western blotting. Three independent replicates are shown.

**C.** Gene Ontology (GO) biological process enrichment analysis of proteins significantly increased abundance in *rad53Δ* compared with WT cells.

**D.** Gene Ontology (GO) biological process enrichment analysis of proteins significantly decreased abundance in *rad53Δ* compared with WT cells.

**E.** Relative protein abundance of Cardiolipin specific deacylase (Cld1) in *rad53Δ* compared with WT cells. Data displayed as means ± SD , n = 3, independent replicates. \*p<0.05, \*\*p<0.01, \*\*\*p<0.001, \*\*\*\*p<0.0001, two-tailed, unpaired Welch's T test.

**F.** Relative protein abundance of Complex I protein (Ndi1), Complex III protein (Qcr7, Cyt1, Qcr2), Complex IV protein (Cox6, Cox10, Cox20, Cox23, Cox26) and Complex V protein (Atp2, Atp4, Atp3, Atp11, Atp23) derived from quantitative proteomic dataset. Data displayed as means ± SD , n = 3, independent biological replicates. \*p<0.05, \*\*p<0.01, \*\*\*p<0.001, \*\*\*\*p<0.0001, two-tailed, unpaired Welch's T test.

**G, H, I.** Heat maps of relative abundances of 'mitochondrial ribosomal small and large subunit proteins', and 'core components of fission-fusion machinery' in *rad53Δ* and WT cells. Heat map showing the median values across three independent biological replicates (n = 3).

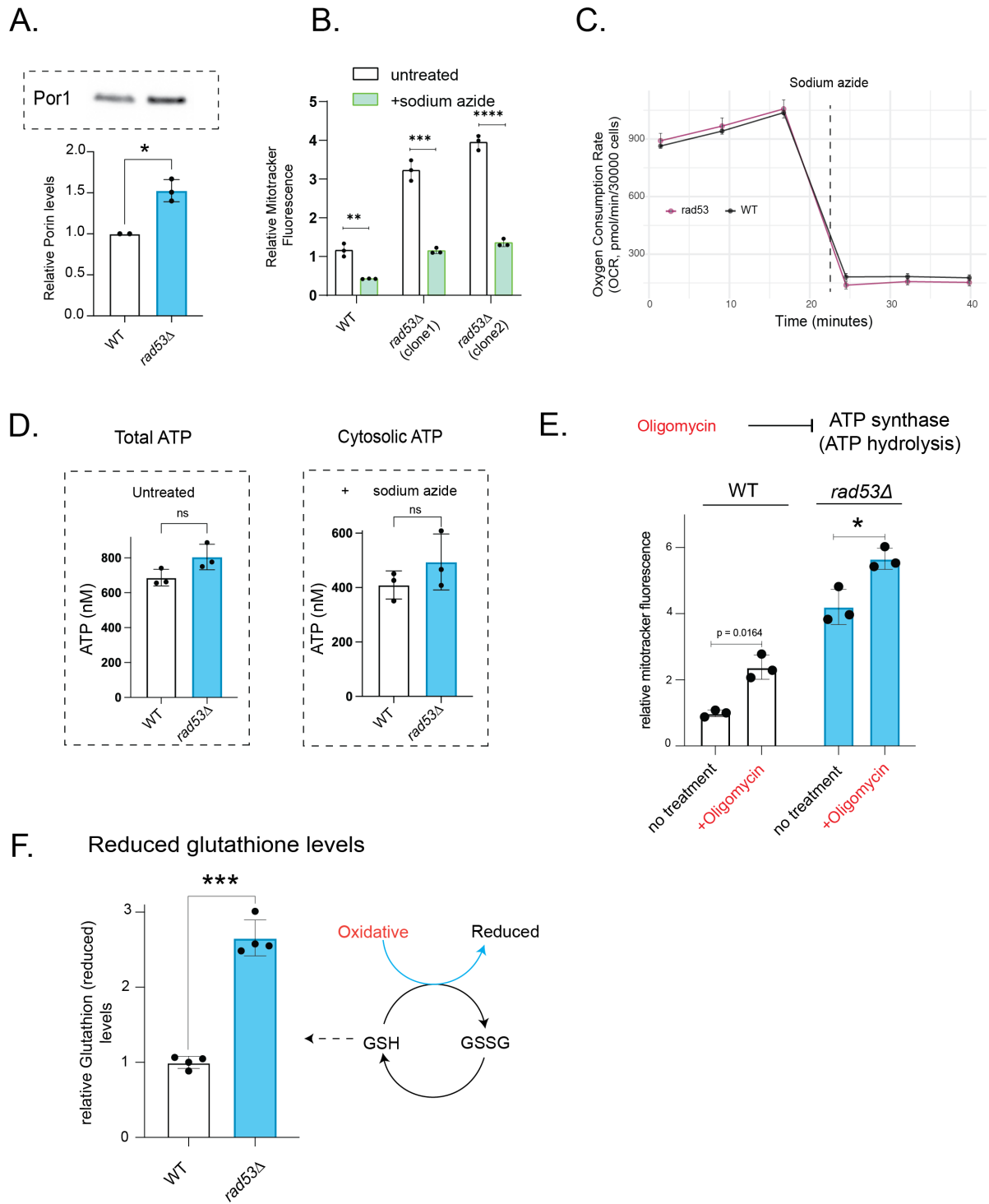

**Fig S4.**

**A.** Representative blot and relative levels of mitochondrial outer membrane porin (POR1) protein levels in *rad53Δ* cells compared to WT cells. Data displayed as means ± SD, $n = 3$. \*$p < 0.05$, two-tailed, unpaired Welch's T test.

**B.** Sodium azide treatment (control) for MitoTracker Red CMXRos staining to quantify mitochondrial potential. Reduction of mitochondrial potential (based on Mitotracker Red intensity) by treating cells with sodium azide (complex IV inhibitor) confirms signal from hyperpolarized mitochondria. Data displayed as means ± SD ,n = 3, independent replicates. \*p<0.05, \*\*p<0.01, \*\*\*p<0.001, \*\*\*\*p<0.0001, two-tailed, unpaired Welch's T test.

**C.** Basal oxygen consumption rate (OCR) of WT and *rad53Δ* cells measured over time using a Seahorse extracellular flux analyzer. Sodium azide treatment served as a negative control for mitochondrial respiration. n = 4

**D.** Concentration of Total ATP and cytosolic ATP. Data displayed as means ± SD ,n = 3, ns denotes non-significant difference, two-tailed, unpaired Welch's T test.

**E.** Relative mitochondrial membrane potential of WT and *rad53Δ* cells following treatment with the ATP synthase inhibitor oligomycin. Oligomycin treatment increased MitoTracker Red CMXRos fluorescence in both WT and *rad53Δ* cells, consistent with inhibition of proton re-entry through ATP synthase. The comparable increase in membrane potential in both strains suggests that the elevated membrane potential in *rad53Δ* cells is unlikely to arise from reverse ATP synthase activity. Data displayed as means ± SD ,n = 3, independent replicates. \*\*p<0.01, \*\*\*p<0.001, two-tailed, unpaired Welch's T test.

**F.** Relative levels of reduced glutathione in WT vs *rad53Δ* cells. Data displayed as means ± SD, n = 4. \*\*\*\*p<0.0001, two-tailed, unpaired Welch's T test

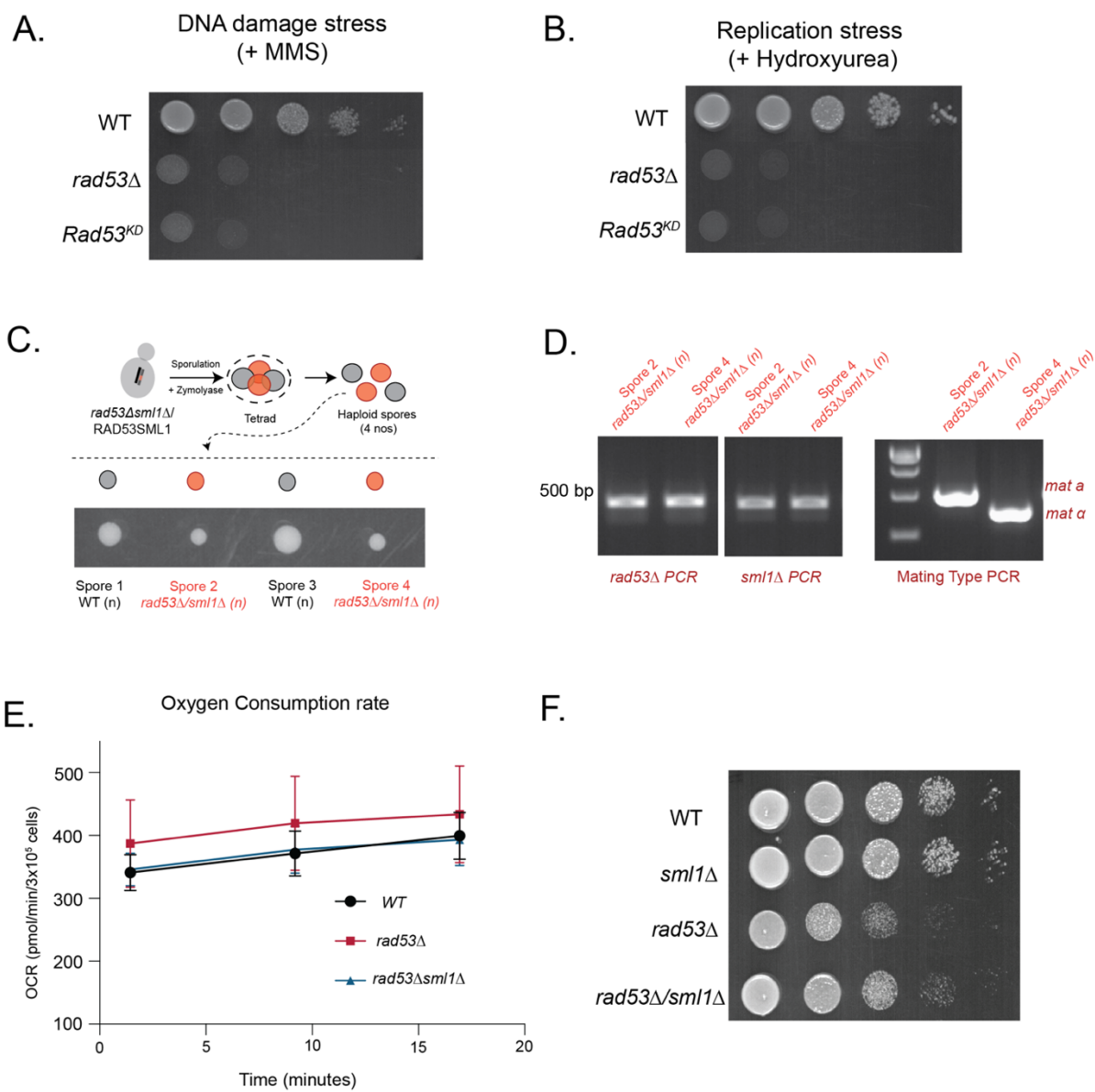

**Fig S5.**

**A.** Growth of WT, *rad53Δ* and Rad53<sup>KD</sup> cells following treatment with methyl methanesulfonate (MMS). Both *rad53Δ* and Rad53<sup>KD</sup> cells exhibit high sensitivity to exogenous DNA damaging agent, consistent with the established checkpoint function of Rad53.

**B.** Growth of WT, *rad53Δ* and Rad53<sup>KD</sup> cells following treatment with hydroxyurea (HU). Both *rad53Δ* and Rad53<sup>KD</sup> cells exhibit high sensitivity to replication stress, consistent with the established checkpoint function of Rad53.

**C.** Viable *rad53Δ/sml1Δ* spores were obtained from diploid parent.

**D.** PCR confirmation of *rad53Δ/sml1Δ* and determination of genotype.

**E.** Basal OCR of *rad53Δ* and *rad53Δ/sml1Δ* cells over time. n = 3

**F.** Growth of *rad53Δ*, *sml1Δ* and *rad53Δ/sml1Δ* cells compared to WT cells. The growth

of *sml1Δ* cells was comparable to that of WT, whereas *rad53Δ* and *rad53Δ/sml1Δ* cells

exhibited reduced growth.

A.

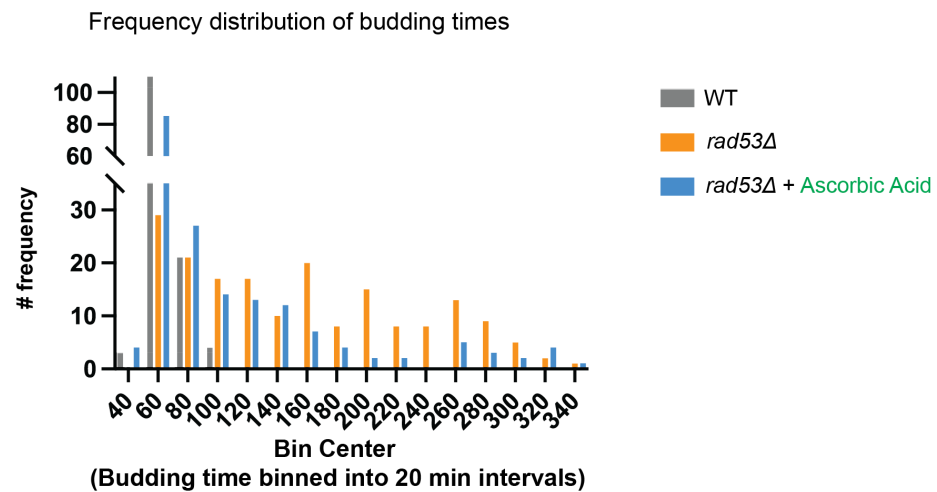

B.

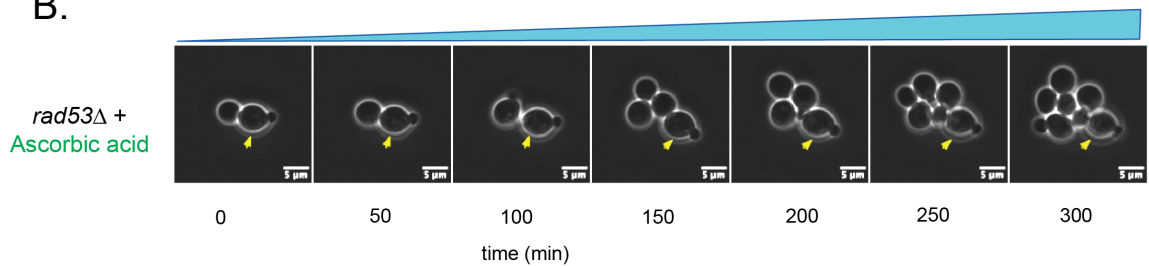

C.

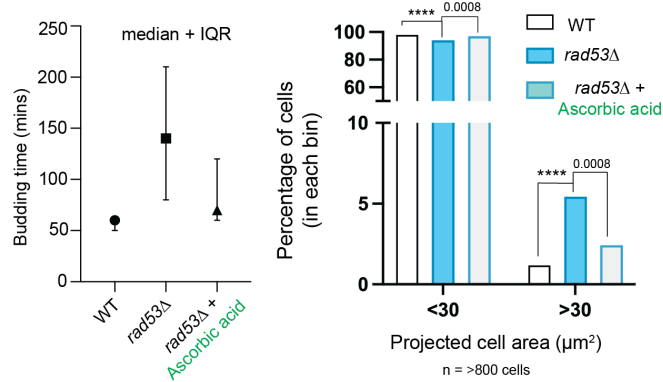

D.

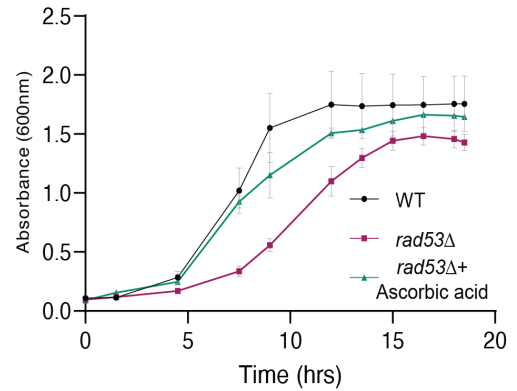

**Fig S6.**

**A.** Frequency distribution of budding times in WT (grey), *rad53Δ* (orange), and ascorbic acid-treated *rad53Δ* cells (blue), with budding times binned into 20-minute intervals. WT cells show a narrow distribution centered around 60 min, whereas *rad53Δ* cells exhibit a broader distribution with a pronounced shift toward longer budding times, indicating increased cell-to-cell variability. Ascorbic acid treatment

restores the budding-time distribution in *rad53Δ* cells, narrowing it predominantly to 60–80 min.

**B.** A representative *rad53Δ* cell (yellow arrow) remains arrested without undergoing cell division throughout the 300-min time course, while neighboring cells continue to divide. Images were acquired at 10-min intervals; representative frames are shown at 50-min intervals. Scale bar, 5 $\mu\text{m}$. Also see, Mov. S6.

**C.** (Left) The median budding time decreased from 140 min (interquartile range, 80– 210 min) to 70 min (interquartile range, 60–120 min), approaching the WT median budding time of 60 min (interquartile range, 50–60 min). (Right) Projected cell areas of WT, *rad53Δ*, and ascorbic acid-treated *rad53Δ* cells, quantified from segmented cell images and grouped into cells with projected areas of either $<30 \mu\text{m}^2$ or $>30 \mu\text{m}^2$. Statistical significance was determined using Fisher's exact test (\*\*\*\*, $P < 0.0001$; WT, $n = 800$; *rad53Δ*, $n = 900$; *rad53Δ* + ascorbic acid, $n = 1000$).

**D.** Growth curve representation of 6D.

#### **Legends for Movies S1 to S6**

**Movie S1 (separate file).** Time-lapse microscopy of WT cells undergoing cell division over 6 h on a YPD agar pad. The movie corresponds to the cell montage shown in the main figure (Fig 1E).

**Movie S2 (separate file).** Time-lapse microscopy of *rad53Δ* cells undergoing cell division over 6 h on a YPD agar pad. The movie corresponds to the cell montage shown in the main figure (Fig 1E).

**Movie S3 (separate file).** Time-lapse microscopy of WT cells undergoing cell division over 6 h on a YPD agar pad. The movie corresponds to the cell montage shown in the main figure (Fig 6C).

**Movie S4 (separate file).** Time-lapse microscopy of *rad53Δ* cells undergoing cell division over 6 h on a YPD agar pad. The movie corresponds to the cell montage shown in the main figure (Fig 6C).

**Movie S5 (separate file).** Time-lapse microscopy of *rad53Δ* cells undergoing cell division over 6 h on a YPD agar pad in the absence of ascorbic acid. The movie corresponds to the cell montage shown in the main figure (Fig 6C).

**Movie S6 (separate file).** Time-lapse microscopy of *rad53Δ* cells on a YPD agar pad supplemented with ascorbic acid over 6 h. The cell indicated by the yellow arrow remains arrested and does not divide during the imaging period.

### **Legends for Datasets S2 to S6**

**Dataset S2 (separate file).** Excel file containing retention times and peak areas for metabolite measurement by LC-MS/MS in WT and *rad53Δ* cells.

**Dataset S3 (separate file).** Excel file containing protein identification and quantitative proteomics data for WT and *rad53Δ* cells, including protein annotations, replicate-level measurements, the number of replicates in which each protein was detected, fold changes, log<sub>2</sub> fold changes, and statistical comparisons between the two conditions.

**Dataset S4 (separate file).** Excel file containing the Gene Ontology (GO) Biological Process enrichment analysis results for upregulated and downregulated proteins, corresponding to Fig. S3C and S3D.

**Dataset S5 (separate file).** Excel file containing the quantitative measurements of selected proteins from the mitochondrial proteome, corresponding to Figs. 3D–F and S3E–I.

**Dataset S6 (separate file).** Excel file containing the quantitative measurements of oxidative stress-related proteins identified in the mitochondrial proteome, corresponding to Fig. 4G.

### **S1 References**

1. Gupta, R. *et al.* A tRNA modification balances carbon and nitrogen metabolism by regulating phosphate homeostasis. *Elife* **8**, (2019).
2. Illuxley, C., Green, E. D. & Dunbam, I. Rapid assessment of *S. cerevisiae* mating type by PCR. *Trends in Genetics* **6**, 236 (1990).

3. Vengayil, V., Niphadkar, S., Adhikary, S., Varahan, S. & Laxman, S. The deubiquitinase Ubp3/Usp10 constrains glucose-mediated mitochondrial repression via phosphate budgeting. *Elife* **12**, (2024).
4. Padovani, F., Mairhörmann, B., Falter-Braun, P., Lengefeld, J. & Schmoller, K. M. Segmentation, tracking and cell cycle analysis of live-cell imaging data with Cell-ACDC. *BMC Biol.* **20**, 174 (2022).
5. Gupta, R., Adhikary, S., Dalpatraj, N. & Laxman, S. An economic demand-based framework for prioritization strategies in response to transient amino acid limitations. *Nat. Commun.* **15**, 7254 (2024).
6. Walvekar, A., Rashida, Z., Maddali, H. & Laxman, S. A versatile LC-MS/MS approach for comprehensive, quantitative analysis of central metabolic pathways. *Wellcome Open Res.* **3**, 122 (2018).
7. Niphadkar, S., Sreedharan, S., Vengayil, V. & Laxman, S. Protocol to quantitatively assess glycolysis and related carbon metabolic fluxes using stable isotope tracing in Crabtree-positive yeasts. *STAR Protoc.* **6**, 103786 (2025).
8. Singh, G., Vengayil, V., Khanna, A., Adhikary, S. & Laxman, S. Active control of mitochondrial network morphology by metabolism-driven redox state. *Proc. Natl. Acad. Sci. U. S. A.* **122**, e2421953122 (2025).
9. Liao, P.-C., Bergamini, C., Fato, R., Pon, L. A. & Pallotti, F. Isolation of mitochondria from cells and tissues. in *Methods in Cell Biology* vol. Volume 155 3–31 (2020).
10. Adhikary, S., Vengayil, V. & Laxman, S. Isolation of Mitochondria From Yeast to Estimate Mitochondrial Pools of Inorganic Phosphate. *Bio. Protoc.* **15**, e5370 (2025).
